# A Bayesian hierarchical analysis of trends and risk to identify regional conservation priorities of Chinstrap Penguins

**DOI:** 10.64898/2026.08.04.742810

**Authors:** Lucas Krüger, Luis Adasme, Carlos Montenegro

**Affiliations:** Instituto Antártico Chileno, Punta Arenas, Chile; Instituto Milénio BASE, Santiago, Chile; Instituto de Fomento Pesquero, Valparaiso, Chile

**Keywords:** Antarctica, Marine Protected Areas, Population Decline, Pygoscelis, Zonation

## Abstract

Chinstrap penguins (*Pygoscelis antarcticus*) in the Antarctic Peninsula and Scotia Arc regions (CCAMLR subareas 48.1 and 48.2) have experienced substantial declines, yet uncertainty remains about regional variation and conservation priorities. We applied a Bayesian hierarchical model to 1,072 nest counts from 194 colonies (1970 to 2024) to quantify population trends, estimate probabilities of exceeding IUCN Red List thresholds, and assess compensation potential across four locations. The regional population declined by 41.9% over three generations, with 91.4% probability of exceeding the 30% Vulnerable threshold but only 9.2% probability of exceeding the 50% Endangered threshold. Declines were spatially heterogeneous: South Shetland Islands (40.2% of regional population) declined by 74.7% with near-certainty of exceeding both thresholds, while South Orkney Islands showed stability with high uncertainty (median -5%, 95% CI: -31.5% to +36.8%). The probability of increasing colonies compensating for regional losses was only 7%. These results support a three-zone conservation strategy for CCAMLR’s proposed Marine Protected Area: urgent protection for the South Shetland Islands, enhanced monitoring for South Orkney and Elephant Islands, and adaptive management for the Antarctic Peninsula. Our uncertainty-quantified framework provides evidence-based guidance for balancing krill fishery access with predator protection in a rapidly changing Southern Ocean.

## Introduction

Southern Ocean Marine Ecosystems have experienced consistent increases in environmental changes, with consequent detrimental effects on biodiversity (Kubiszewski et al., 2025; Robinson, 2022). Of particular importance are land-based marine predators, which face impacts from environmental change at both land and sea habitats (Blondin et al., 2022; Krüger, 2022).

In the last decade, climate-induced redistribution of the main primary consumer (Atkinson et al., 2022; Kawaguchi et al., 2023), effects on breeding habitat (Descamps et al., 2023; Gimeno et al., 2025; Machado-Gaye et al., 2025; Youngflesh et al., 2020), and increases in the concentration of commercial fisheries (Grémillet et al., 2018; Krüger et al., 2021; Watters et al., 2020) have all been recorded. Consequently, dramatic changes in population numbers have been observed throughout the Southern Ocean across different levels of marine food webs. Predator populations (or meta-populations) previously assumed stable or recovering from past impacts have suffered substantial declines throughout their range in the Antarctic Peninsula and adjacent islands (Braun et al., 2021; Fretwell et al., 2023; Krause et al., 2024; Petry et al., 2018).

The chinstrap penguin (*Pygoscelis antarcticus*) has suffered steep declines throughout its range, particularly in the subareas 48.1 and 48.2 (Antarctic Peninsula, South Shetland Islands, and South Orkney Islands; Strycker et al., 2020; Talis et al., 2023). That is a region that holds between 50% and 60% of the global population (Krüger, 2023; Strycker et al., 2020). There, colonies have had declined over 30% in 30 years (Krüger, 2023; Oosthuizen et al., 2024), and some of the largest colonies have declined by more than 50% over 40 years (Hinke et al., 2025; Pizarro et al., 2026; Talis et al., 2023).

Globally, the species is considered moderately depleted, partly because stable or increasing colonies in the South Sandwich Islands appear to compensate for widespread losses elsewhere. However, this assessment carries high uncertainty due to the low number of samples, low quality of counts, and absence of a historical baseline in the South Sandwich Islands (Hart and Convey, 2018; Lynch et al., 2016; Strycker and Lynch, 2022).

These regional declines occur within the Commission for the Conservation of Antarctic Marine Living Resources (CCAMLR) Domain 1, where CCAMLR is actively evaluating a Marine Protected Area (MPA) proposition (Brooks et al., 2020; Sylvester and Brooks, 2019). CCAMLR operates under a dual mandate: to allow rational use of marine living resources, notably the Antarctic krill fishery, while applying precautionary protection to vulnerable ecosystem components (Graham and Webb, 2022; Guggisberg, 2025) including krill-dependent predators such as the chinstrap penguin (Krüger et al., 2024).

The proposed MPA would require zoning that balances both objectives, yet such zoning depends on knowing where Chinstrap penguin colonies are declining most rapidly and where data gaps prevent reliable assessment. In subareas 48.1 and 48.2, Chinstrap penguin colonies have experienced steep declines and show high probabilities of crossing IUCN Red List thresholds. Therefore, identifying priority zones for strict protection (e.g., foraging areas around rapidly declining colonies) versus zones needing improved long-term monitoring (e.g., under-sampled or apparently stable colonies) is crucial for an evidence-based MPA design. Without regionally explicit, uncertainty-quantified trends, CCAMLR risks either under-protecting collapsing populations or unnecessarily restricting fisheries where impacts are minimal.

This study addresses these conservation and management needs using a Bayesian hierarchical model applied to all available chinstrap penguin nest counts from subareas 48.1 and 48.2 (1970 to 2024). The approach explicitly accounts for variable survey accuracy, colony-specific random effects, and spatial variation among four locations: Antarctic Peninsula, South Shetland Islands, Elephant Islands, and South Orkney Islands. Were quantified: (i) local and regional percent change over three generations (28.2 years), (ii) the probability of exceeding IUCN decline thresholds (30% and 50%), and (iii) the capacity of increasing colonies to compensate for declining ones. Finally, population-weighted posterior aggregation were used to propose regional conservation priorities, distinguishing areas where protection is urgently needed from areas where enhanced monitoring should be prioritised. Results provide a robust, uncertainty-quantified basis for CCAMLR’s MPA zoning and adaptive management of the krill fishery in Domain 1.

## Methods

### Data Preparation and Colony Classification

Chinstrap penguin nest counts data was downloaded from the MAPPPD database (version 4.4; (Humphries et al., 2017; Lynch et al., 2026) for subareas 48.1 and 48.2. MAPPPD compiles all publicly available penguin count data from Antarctica. Following Croxall and Kirkwood (1979), each count record in MAPPPD includes a quality flag from 1 to 5 indicating survey accuracy. Quality flags of 1 represent the highest accuracy (e.g., ground counts covering entire breeding sites with ∼ 5% to 10% error), while flags of 5 represent the lowest accuracy (e.g., estimates derived from satellite imagery or modelled output). All records were included in the analysis and accuracy was incorporated as a random effect in the hierarchical model to account for potential systematic differences in baseline counts and trends among accuracy levels. The analysis was restricted to surveys conducted from 1970 onwards from colonies with a minimum of two counts in different years. A total of 1,072 counts of 194 colonies were available. Colonies were grouped by location as Antarctic Peninsula (AP), South Shetland Islands (SSI), Elephant Islands (EI) and South Orkney Islands (SOI); colonies were also grouped based on their mean colony size (mean number of nests) using a gap analysis (Supplementary File 1 fig S1 and S2) which grouped them in Small (up to 550 nests), medium (over 550 up to 10,000 nests) and large (over 10,000 nests).

#### Bayesian Hierarchical Model Structure

A Bayesian hierarchical model (BHM) was applied using the `brms` package (Bürkner 2017, Carpenter et al. 2017) in R (R Core Team 2024). BHMs are a robust approach for dealing with data-poor groups by “borrowing strength” from data-rich sites within the nested structure of the data (White et al., 2024; Xu et al., 2020). This approach is therefore well-suited for analyzing counts of penguin nests throughout Antarctica, as there is a mix of colonies: some have been counted only twice, many have been counted 5 to 15 times, and some have more than 15 counts.

Bayesian inference was implemented using Markov Chain Monte Carlo (MCMC) sampling. Four independent chains were run for 4,000 iterations each, discarding the first 1,000 as warmup, yielding 12,000 posterior draws for inference, which generates a representative sample of from the joint posterior distribution of all model parameters (Hoffman & Gelman 2014). Each draw is a single set of parameter values consistent with both the observed data and specified prior distributions. Collectively, the draws approximate the full posterior distribution, allowing for comprehensive uncertainty quantification without relying on asymptotic approximations. For derived variables, such as percent change over three generations, population-weighted trends, and probabilities of exceeding IUCN decline thresholds, the relevant transformations to each posterior draw were applied individually. This approach, known as posterior propagation of uncertainty (Candela et al., 2003; Draper, 1995), ensures that all sources of uncertainty are fully represented in final estimates. Summaries such as posterior medians, 95% credible intervals, and probability statements (i.e., P[decline ≥30%]) were then calculated directly from the empirical distribution of these transformed draws.

The model used a student-t distribution normalized (centered) around the variance with value-dependent degrees of freedom (smaller values mean heavier tails) of the expected mean nest counts (log-transformed to normalize residuals) for each site varying to the survey year centered at the mean (1997), Centering the years improve MCMC sampling efficiency and allow interpretation of the intercept as the estimated log-count at the mean year. Site-specific random intercepts and slopes were used, capturing the effect of deviations from the population-level trend for each colony over the general trend, and also random effects for accuracy level (1 to 5) were used, accounting for systematic differences in survey methodology. See detailed model specification in Supplementary File 2.

The inclusion of both site-specific random slopes and location interactions allows the model (Simpson et al., 2017)to partition variance into: (1) domain-1 level trends, (2) local trends, (3) colony-level variation around these trends, and (4) measurement-related variation.

Weakly informative priors (Supplementary File 2) were applied to regularize parameter estimates (Gelman et al., 2017). The exponential prior for variance components was chosen following (Simpson et al., 2017), who recommend regularizing priors for hierarchical models to prevent overfitting.

Following a Student-t distribution to accommodate heavy-tailed residuals, a weakly informative Gamma prior was placed on the degrees of freedom parameter. This prior encourages moderate to heavy tails while allowing the data to inform the exact shape (with a prior mode approximately Gaussian) and sufficient variance to allow degrees of freedom to fall as low as 1 (Cauchy) if supported by the data (Juárez and Steel, 2010; Villa and Walker, 2014).

Convergence was assessed using the scale reduction factor (R-hat; Gelman and Rubin, 1992), requiring R-hat ≤ 1.01 for all parameters (Vehtari et al., 2021). Effective sample sizes (ESS) were checked to ensure sufficient posterior information (all ESS > 1000).

Model fit was assessed using leave-one-out cross-validation LOO with moment matching (Vehtari et al., 2017) and using the Bayesian R^2^ (Gelman et al., 2019). Only 0.01% of draws represented divergent transitions. . The LOO validation identified 26 observations (2.5%) with k > 0.7, indicating some influence, but moment matching provided robust estimates. Given the convergence values (all R-hat = 1.00), high bayes R^2^ (0.946 [0.937, 0.954]), large effective sample sizes and model predictive checks (see below) the model was considered adequate.

Model adequacy was evaluated through posterior predictive checks (Gelman et al., 2013). A thousand replicated datasets were generated from the posterior predictive distribution and compared to observed data using kernel density overlays. Systematic discrepancies between observed and replicated data would indicate model misspecification.

### Derivation of Effect Sizes and Decline Probabilities

To facilitate biological interpretation, model parameters were transformed to percent change over three generations (28.2 years): %Δ = [exp(β × 28.2) - 1] × 100, where β represents either global trend or location-specific trend coefficients. Uncertainty was propagated by applying this transformation to all posterior draws, yielding full posterior distributions of percent change.

Median percent change with 50% and 95% credible intervals, probability of exceeding IUCN Red List thresholds, P(decline ≥30%) and P(decline ≥50%) were calculated for each location.

### Regional-Level Inference

To estimate regional population trajectory accounting for colony sizes, a weighted posterior aggregation approach was implemented: The long-term mean count (N_i_) and population weight for each colony *i* was calcualted: w_i_ = N_i_ / ΣN_i_ ; the population-weighted trend was calculated for each posterior draw *d*: β_pop(d)_ = Σ w_i_ × β_i_ (d), where β _i_ (d) is the site-specific slope for colony *i* in draw *d*. The posterior distribution of regional population percent change was derived as above.

Compensation effects were further quantified by calculating, for each posterior draw: total expected increase: Σ N_i_ × [exp(β _i_ × 28.2) - 1] for sites with β _i_ > 0; total expected decline: Σ |N_i_ × [exp(β _i_ × 28.2) - 1]| for sites with β _i_ < 0; net change = increase - decline; compensation probability = P(net change > 0).

To assess population trends relative to IUCN Red List criteria, estimated annual rates of change forward over three generations were projected (28.2 years). This time frame is a fixed benchmark (number of generations) for risk assessment and does not reflect the actual duration of the monitoring data, which spanned 54 years (1970–2024). For each posterior draw from the Bayesian model, the expected percent change over 28.2 years was calculated as %Δ = [exp(β × 28.2) − 1] × 100, where β represents the estimated annual log-scale trend (either global or location-specific). This approach assumes that the average annual rate of change estimated from the historical period remains constant over the projection interval, an assumption that allows consistent comparison across species and populations. However, this represents a projection of past trends rather than a forecast incorporating potential future environmental changes. The full uncertainty from the model is propagated through this calculation by applying the transformation to all posterior draws, yielding posterior distributions of percent change that reflect both estimation uncertainty and, through the site-level random effects, heterogeneity among colonies. Probabilities of exceeding IUCN decline thresholds (30% and 50%) were then computed directly from these posterior distributions.

### Data Availability and Reproducibility

Detailed R-code is available in Supplementary File 3, and alternatively in (Krüger et al., 2026). Data is openly available at (Krüger et al., 2026: https://doi.org/10.5281/ZENODO.21826915) whose use needs to refer to penguinmap.com (Humphries et al., 2017; Lynch et al., 2026). Antarctic shapefiles used for maps are from (Gerrish et al., 2026) and statistical subareas shapefiles from CCAMLR GIS (https://gis.ccamlr.org/).

## Results

### Model Diagnostics

The Bayesian hierarchical model showed excellent convergence (all R-hat = 1.00) and high effective sample sizes (minimum Bulk-ESS = 2,377, minimum Tail-ESS = 3,995), indicating reliable posterior sampling (fig S3-S4). Bayesian R^2^ indicated that the model explained substantial variation in log-transformed counts (0.946 [0.937, 0.954]). Leave-one-out (LOO) cross-validation with moment matching found 1 divergent transitions (0.01% of draws) and 26 observations (2.4%) with k > 0.7.

### Survey Accuracy and Trend Estimates

The model revealed that measurement quality slightly affected baseline counts (intercepts) but not trend estimates (slopes). Slight variations in the intercept of accuracy class 3 and 4 were found, but all error bars crossed zero (fig S5a). Accuracy random slopes were very low with negligible error bars (fig S5b). Correlation between site-level random slope and random intercept was weak (0.22 [IQR 95% 0.06 to 0.37]).

### Local scale trends

Larger colonies were located in the South Shetland Islands and Elephant islands, where also the declines predominantly occurred (figure 1). The Antarctic Peninsula colonies are mostly stable or slightly increasing (figure 1). Elephant Islands and South Orkneys Islands have, together with South Shetland Islands, an important number of large colonies, but while there is a balance in number of decreasing and increasing colonies in South Orkneys, In South Shetland and Elephant Islands diminishing colonies are the norm (figure 1).

**Figure 1.**
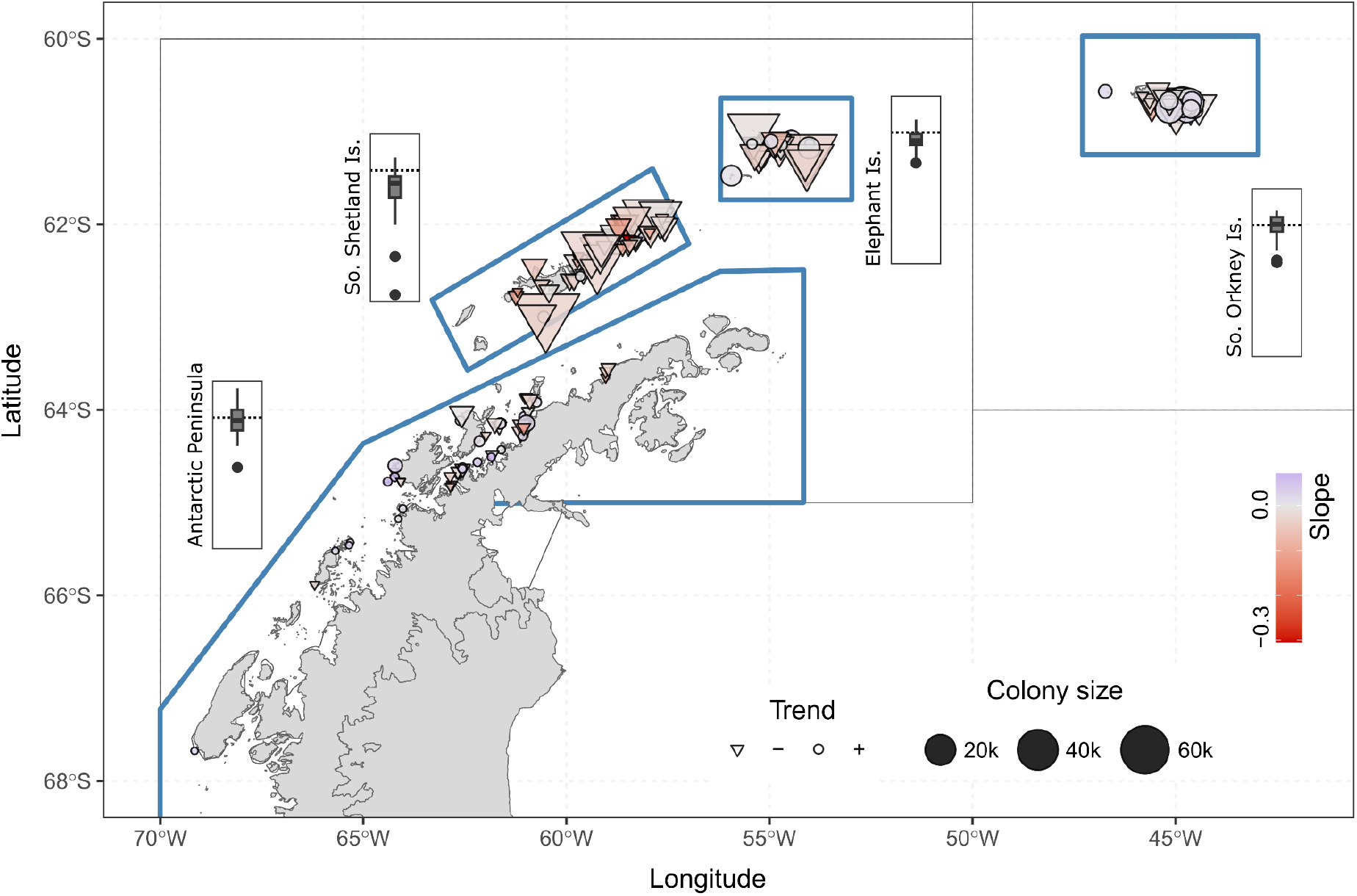
Chinstrap penguin (*Pygoscelis antarcticus*) colonies in the Domain 1 and their trend (site-level slope) calculated from the site-effect of a Bayesian mixed model. Positions of decreasing colonies (-) are represented by the inverted triangle and stable or increasing colonies (+) are represented by circles. Symbols are sized based on mean colony size (number of nests) and colored by the site-level slope. Boxplots show, for each location, the frequency distribution of slope values in 25% quantiles around the median (vertical black line) and outliers (black dots).

### Risks

The overall change in number of nests was -41.9% in three generations, with over 90% chance of declining by 30% in three generations, but a low chance (9.2%) of declining 50% in three generations (Table 1). The probability of increasing colonies compensating for the declining ones was very low, and the overall decline in three generations remained in the hundreds of thousands nests lost in three generations (table 1).

**Table 1.** Regional chinstrap penguin (*Pygoscelis antarcticus*) population trend, risk of crossing 30% and 50% decline thresholds within three generations,probability that increasing colonies compensate for declining ones, and the median net population change. Median values and Inter-Quartil Range IQR within brackets.

| Metric | Estimate [IQR] |
| --- | --- |
| Population trend (% change) | -41.9% [-56.1% to -17.8%] |
| Risk of $\geq 30\%$ decline | 91.4% [90.9% to 91.9%] |
| Risk of $\geq 50\%$ decline | 9.2% [8.7% to 9.8%] |
| Probability of compensation | 7% [6.5% to 7.5%] |
| Net population change (thousand nests) | -211.5 [-391.5 to 145.4] |

South Shetland and Elephant Islands colonies together had almost 70% of the regional number of nests, but with disparity in the levels of decrease experienced by the colonies, ∼75% and ∼39% decreases, respectively (Table 2). South Shetland Island colonies were under certainty of reaching 30% and 50% declines over three generations, while Elephant Island colonies were under a high risk of 30% decline (Table 2). The Antarctic Peninsula median trend was negative, but the risk that the decline would cross the 30% or 50% was low (Table 2). On the other hand, South Orkney Island median change was close to zero, but with wide credible intervals, suggesting there are differences in colony-level trends within the location (table 2).

**Table 2.** Local chinstrap penguins (*Pygoscelis antarcticus*) contribution to the number of nests for the regional population, the median local population trend and risk of crossing 30% and 50% decline thresholds within three generations.

| location | Population share | Median % change [95% CI] | Risk of ≥30% decline | Risk of ≥50% decline |
| --- | --- | --- | --- | --- |
| Antarctic Peninsula | 2.3% | -20.6% [-40.5% to 15.7%] | 15.5% | 0.7% |
| South Shetland Islands | 40.2% | -74.7% [-81.1% to -62.5%] | 99.8% | 99.4% |
| Elephant Islands | 29.3% | -38.8% [-55.5% to -16.3%] | 86.2% | 7.0% |
| South Orkney Islands | 28.2% | -5% [-31.5% to 36.8%] | 3.1% | 0.4% |

### Counts and colony size

Locations with better coverage are the Antarctic Peninsula and South Shetland Islands, which have on average 7 to 10 counts per colony, but only for small and medium colonies (Figure 2a). Large colonies have on average two or three counts, although they are less numerous than medium and small colonies (figure 2a). There is no obvious relation of colony size and trend, although there is a slight geographical variation suggesting that larger colonies further north are decreasing more than the smaller colonies in the Antarctic Peninsula, with the exception of South Orkney Islands (Fig 2 b). Within location variability also suggests geographical gradients in the Antarctic Peninsula and South Shetland Islands (Fig 2b).

**Figure 2.**
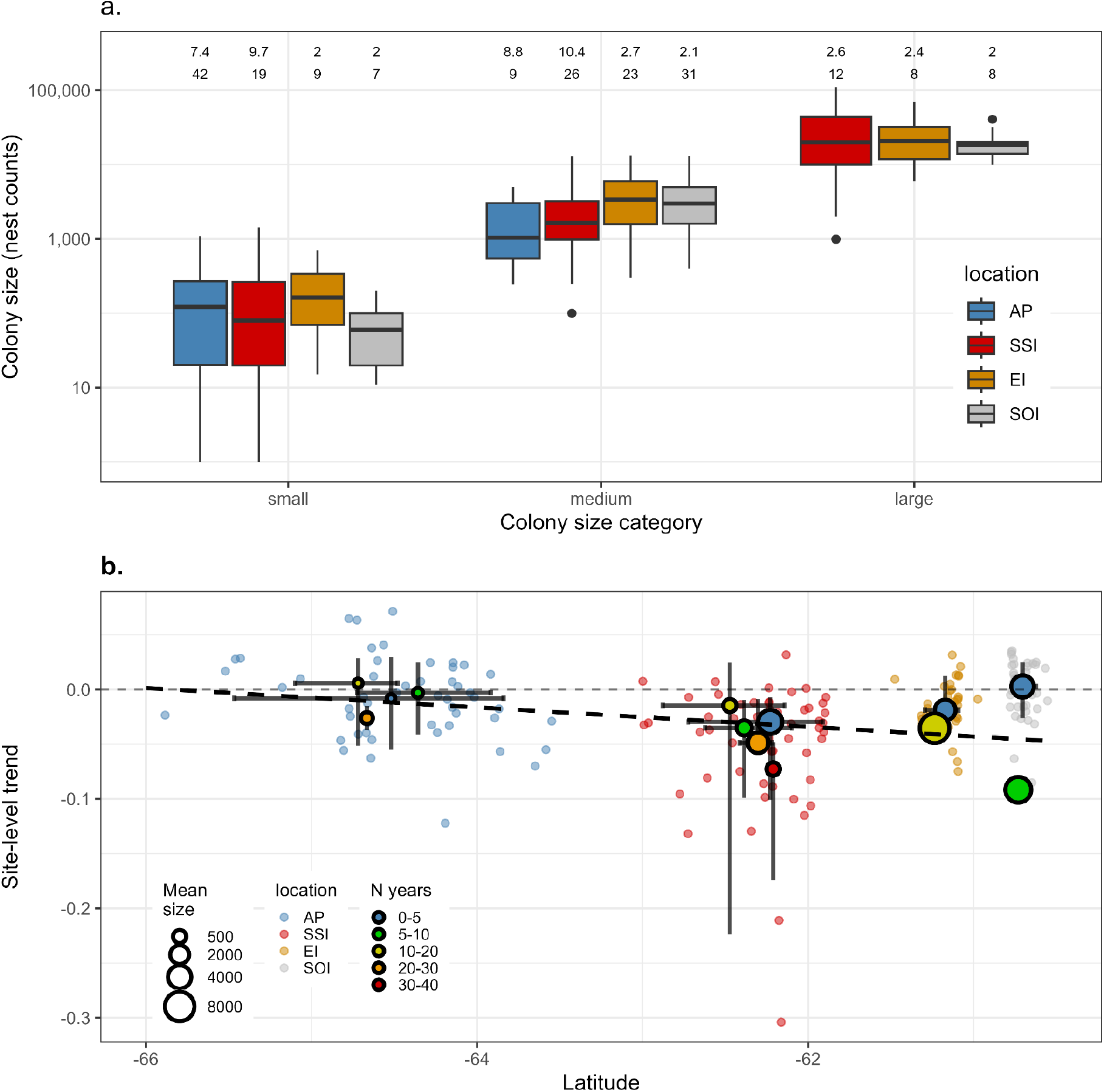
Chinstrap penguin (*Pygoscelis antarcticu*s): (a) colony size by location, showing the mean number of counts (top numbers) and number of colonies (bottom numbers); and (b) latitudinal variation in site-level trends (slopes) for each colony, with points colored by location and centroids ± 75% IQR shown for colonies grouped by the number of years with count or nest-estimation data. Locations in both panels are the Antarctic Peninsula (AP), South Shetland Islands (SSI), Elephant Island (EI), and South Orkney Islands (SOI). See Figure 1 for geographical boundaries of these locations.

## Discussion

### Regional Declines and Conservation Priorities

This study provides a comprehensive Bayesian hierarchical analysis of Chinstrap penguin population trends across subareas 48.1 and 48.2, quantifying both the magnitude of declines and the uncertainty surrounding these estimates. Results revealed a regional population decline of 41.9% over three generations, with a 91.4% probability of exceeding the IUCN Red List threshold for a vulnerable classification (30% decline over three generations). However, the probability of crossing the more severe 50% decline threshold remains low (9.2%), suggesting that while the species is moderately depleted regionally (matching findings in (Strycker and Lynch, 2022), it has not yet reached the threshold for Endangered status in Domain 1.

These regional trends mask substantial spatial heterogeneity that carries direct implications for MPA design. The South Shetland Islands hold 40.2% of the regional population and have experienced catastrophic declines of 74.7% (95% CI: 81.1% to 62.5%), with near-certain probabilities of exceeding both the 30% (99.8%) and 50% (99.4%) decline thresholds. Elephant Islands, holding an additional 29.3% of the population, show severe declines of 38.8% with an 86.2% probability of exceeding 30% decline. Those are consistent with levels of decline found by (Talis et al., 2023). In contrast, the Antarctic Peninsula (2.3% of population) showed moderate declines with high uncertainty (median -20.6%, 95% CI: -40.5% to +15.7%), while South Orkney Islands (28.2% of population) exhibit relative stability (median -5%, 95% CI: -31.5% to +36.8%). These divergent trajectories indicate that regional stability or recovery would be driven primarily by the South Orkney and Antarctic Peninsula populations, but their contribution to total population size (30.5% combined) make them incapable of compensating for losses in the South Shetland and Elephant Islands.

A key finding with direct management relevance is the low probability of compensation, it is the chance that increasing colonies could offset declines elsewhere. The population-weighted analysis indicates that even if every increasing colony realized its full positive trajectory, the net change over three generations would still represent a loss of over 200,000 nests (95% CI: -391,500 to +145,400). The wide credible interval for net change reflects uncertainty about whether the few increasing colonies, predominantly small colonies on the Antarctic Peninsula and South Orkney Islands, can sustain their positive trends. However, the posterior probability of any net increase is only 7%, providing evidence against meaningful compensation.

Important to notice, this result does not take into account colonies on subareas 48.4 (South Sandwich Islands) which are known to be among the largest chinstrap colonies with most of them being stable or increasing (Lynch et al., 2016; Oosthuizen et al., 2024; Strycker et al., 2020). On the other hand, those estimations are from over 10 years ago, and recently updated data indicate a recent ⅓ decline in this area (Gregory and Belchier, 2026; Ratcliffe et al., 2025), what would bring the global trend of the species to an Endangered status (Ratcliffe et al. in press).

### Data Gaps and Monitoring Priorities

Our analysis reveals critical data deficiencies that constrain inference and should guide future monitoring investments. Large colonies (>10,000 nests), which disproportionately contribute to population trends, have on average only 2 to 3 counts across the 54-year study period, compared to 7 to 10 counts for small and medium colonies. This disparity is particularly concerning because the largest declines have occurred precisely in these large colonies, predominantly in the South Shetland and Elephant Islands. The South Shetland Islands have been sampled relatively consistently, but almost exclusively for small and medium colonies that are not representative of the regional population trajectory. Elephant Islands and South Orkney Islands contain numerous large colonies, but only a handful have sufficient temporal replication to support robust trend estimation. The clear mismatch between monitoring effort and the demographic contribution of individual colonies increases uncertainty in the estimated trends and limits our ability to test for environmental effects on the colonies that drive regional population dynamics. A more focused sampling strategy, one that is representative of the regional population structure, is therefore needed.

From a management perspective, these data gaps suggest two priority actions. First, targeted resampling of large colonies in the South Shetland and Elephant Islands should be urgently prioritized to reduce uncertainty in decline estimates and improve early warning of accelerating losses. Second, the South Orkney Islands, where large colonies appear relatively stable but have extremely low sampling intensity, should be designated as a sentinel monitoring region. The wide credible interval around the South Orkney trend estimate (-31.5% to +36.8%) reflects the uncertainty about whether this population is stable, declining, or increasing. Resolving this uncertainty is essential for understanding whether the South Orkney population could serve as a future refuge or is already experiencing cryptic declines.

### Implications for CCAMLR MPA Design

The proposed MPA in CCAMLR Domain 1 requires zoning that balances krill fishery access with protection of krill-dependent predators (Krüger et al., 2024). This study provides quantitative, uncertainty-quantified evidence to inform this zoning process taking into account the Chinstrap Penguins. Three distinct management zones can be proposed:

Zone 1, Urgent Protection (South Shetland Islands). Colonies in this region have declined by nearly 75% over three generations, with 99.8% probability of exceeding the 30% decline threshold. The probability of crossing 50% decline is similarly 99.4%, indicating that without intervention, this population is on track for potential extirpation. We recommend that foraging areas within a 50-100 km radius of remaining large colonies in the South Shetland Islands receive strict protection from krill fishing, following the precautionary approach required under CCAMLR’s mandate(Graham and Webb, 2022). Given that this region holds 40% of the regional population, its protection should be the highest priority. Understanding which periods of the breeding cycle and which life stages might have been impacted by krill fishery also is a research priority for the South Shetlands (Hinke et al., 2020; Krüger et al., 2021; Watters et al., 2020). Understanding the mechanisms of potential impact will allow for a more efficient zonation of fishery closures to reduce potential impacts, both spatially and temporally (Krüger et al., 2024). For instance, krill fishery industry have limited their activity near large penguin colonies during critical breeding stages of penguins (Godø and Trathan, 2022), but those do not cover the pre- and pos-moulting period for Chinstrap Penguin, that can extend into late April (Black et al., 2018), and might as well be under ongoing changes (Juarez Martinez et al., 2026). Additionally, the South Shetland Islands also sustain most of the Antarctic Fur Seal colonies in the area, which are genetically diverse and have also experienced declines as severe as the chinstrap penguins (Krause et al., 2024, 2022).

Zone 2, Enhanced Monitoring (South Orkney and Elephant Islands). The Elephant Islands population has declined severely (38.8% median decline) but with greater uncertainty, while the South Orkney population appears stable but with extremely wide credible intervals. For Elephant Islands, we recommend moderate fishing restrictions combined with targeted annual monitoring to determine whether declines are accelerating or stabilizing. Studies published in the area provide a solid baseline, but most of the colonies still lack a consistent temporally regular sampling (Petry et al., 2018; Strycker et al., 2021). For South Orkney Islands, we recommend prioritized investment in high-frequency monitoring (e.g., annual ground or drone surveys of large colonies) to detect emerging declines before they become irreversible.

Zone 3, Adaptive Management (Antarctic Peninsula). The Antarctic Peninsula contains mostly small colonies with moderate declines. This region has high sampling intensity, yet its colonies contribute only 2.3% to the regional population. On the other hand, sectors of the Peninsula, such as Gerlache Straight and Joinville Island, are of high importance for Krill recruitment stages (Krüger et al., 2024; Meyer et al., 2023) and baleen whales throughout summer and winter (Bahlburg et al., 2025; Krüger et al., 2024).

It is also important to highlight that that are important gaps in the northern sector of the AP, in the Bransfield Straight, where some colonies of medium to large size are known (i.e. Tupinier Islands or Lafarge Rocks) which have only one count. This sector represents the larger data-gap on both subareas at the same time that it is closer to the most fished hotspot in the subarea 48.1 (Santa Cruz et al., 2022, 2018). We recommend an adaptive feedback management approach (i.e. Klein and Watters, 2020) in the Southern Bransfield Strait, where krill fishing is permitted but subject to regular review based on regular data updates.

### Comparison with Previous Assessments

The estimate of a 41.9% regional decline over 28.2 years is broadly consistent with previous studies that reported declines exceeding 30% over similar timeframes (Krüger, 2023; Oosthuizen et al., 2024; Strycker et al., 2020), and with newly updates on large colonies (Gregory and Belchier, 2026; Hinke et al., 2025; Pizarro et al., 2026). However, this study advances previous work in three important ways. First, by explicitly modeling uncertainty through a Bayesian hierarchical framework, we provide probabilistic statements about management-relevant thresholds (e.g., P(decline ≥30%) = 91.4%) rather than point estimates with credible intervals. This probabilistic framework aligns directly with the precautionary approach required under CCAMLR’s decision-making processes. Second, our population-weighted aggregation reveals that the magnitude of decline in large colonies, often under-sampled, is substantially greater than would be inferred from unweighted colony averages. Third, our quantification of compensation probability (7.0%) provides formal evidence against the hypothesis that increasing colonies can offset regional losses, a question that previous studies have addressed only qualitatively.

### Uncertainties and Limitations

Some limitations warrant consideration. First, projection of percent change over three generations assumes that historical trends continue linearly into the future. This assumption is one of the standard approaches for extrapolating population trends under IUCN Criterion A (Standards and Petitions Committee of the IUCN Species Survival Commission, 2024), but it has also been identified as a potential source of bias when the true pattern of decline is non-linear or when threat processes are accelerating (Akçakaya et al., 2006). Second, our analysis is restricted to subareas 48.1 and 48.2, which hold 50% to 60% of the global population, but we cannot extrapolate to the South Sandwich Islands, where stable or increasing colonies may partially offset the declines documented here. However, given the high uncertainty surrounding South Sandwich Islands estimates (Hart and Convey, 2018; Lynch et al., 2016), and updated data potentially revealing decreases (Gregory and Belchier, 2026; Ratcliffe et al., 2025), we echo previous calls for targeted surveys in that region to establish a reliable baseline.

Finally, our model does not explicitly incorporate environmental covariates (such as sea-ice extent, krill biomass, or sea surface temperature) nor does it include fishery catch data. Although this was an intentional choice aimed at estimating trends and risks rather than attributing causation, it also means we cannot determine whether observed declines are driven by climate change, fishery competition, or their interaction. Understanding the relative contribution of the fishery to these declines, and consequently the magnitude of the population’s response to the proposed spatial protection, has direct implications for fisheries management and for assessing the effectiveness of the proposed zonation. This can be achieved through the implementation of a robust monitoring program that supports an adaptive feedback management strategy, allowing for the verification of zonation performance and the adjustment of conservation measures as needed.

## Conclusions

Chinstrap penguins in subareas 48.1 and 48.2 have declined by approximately 42% over three generations, with a 91% probability of meeting IUCN criteria for a Vulnerable classification. These declines are not evenly distributed: the South Shetland Islands (40% of the regional population) have experienced catastrophic losses approaching 75%, while the South Orkney population remains stable but highly uncertain. The probability that increasing colonies could compensate for declines is only 7%, meaning that regional recovery is unlikely without active intervention. For CCAMLR’s proposed MPA in Domain 1, these results support a three-zone strategy: urgent protection for the South Shetland Islands, enhanced monitoring for South Orkney and Elephant Islands, and adaptive management for the Bransfield Strait. The Bayesian framework developed here provides a template for uncertainty-quantified, management-relevant assessments that can be updated iteratively as new data become available, an essential capability for adaptive management in rapidly changing Southern Ocean ecosystems.

## Acknowledgements

This study was supported by the Marine Protected Areas Program of Instituto Antártico Chileno (AMP 24 09 052), and by ANID - Millennium Science Initiative Program - ICN 2021_002 (BASE).

