## Supplementary File 1 for "A Bayesian hierarchical analysis of trends and risk to identify regional conservation priorities of Chinstrap Penguins"

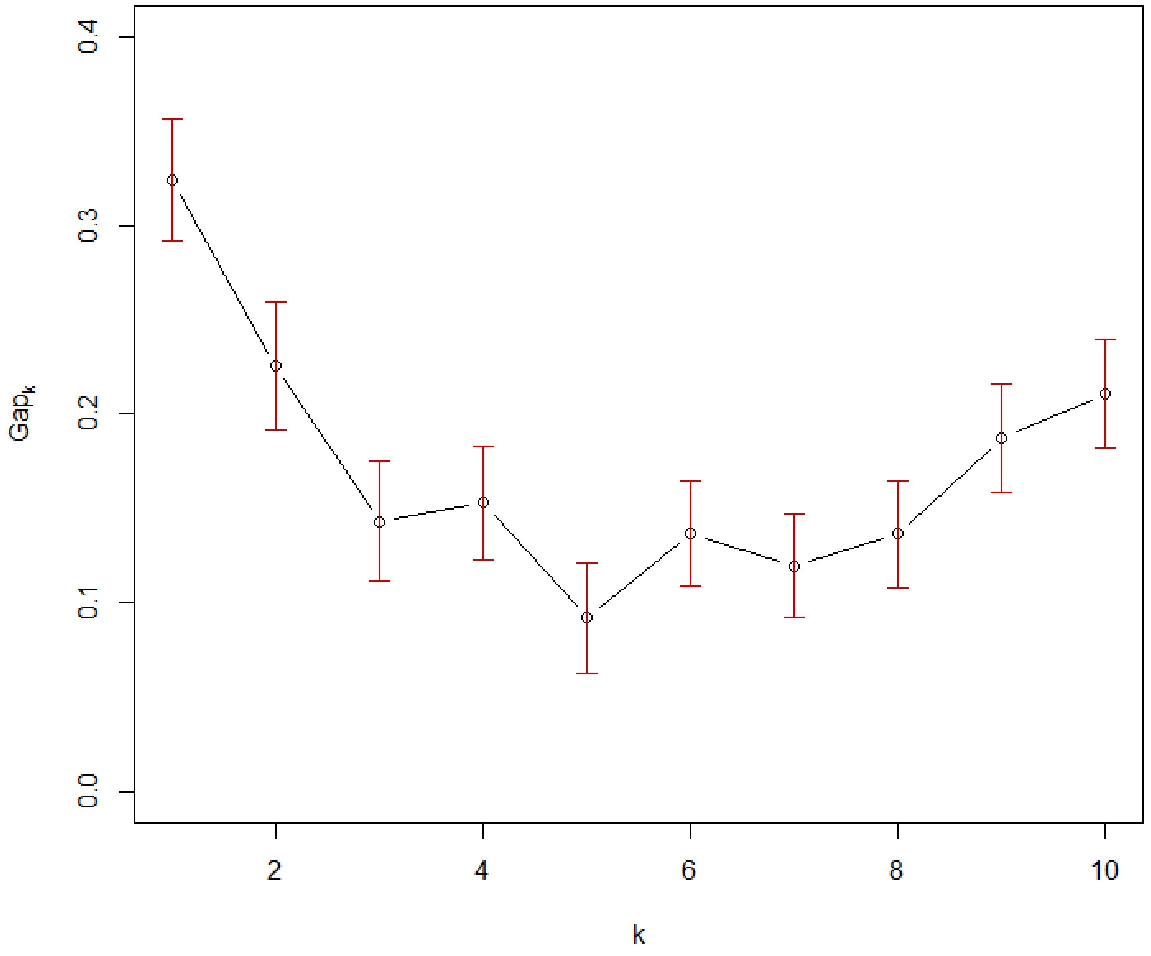


Fig S1. Gap statistic values for chinstrap penguin (*Pygoscelis antarcticus*) colony size clustering. The plot shows the gap statistic ($Gap_k$) as a function of the number of clusters (k) ranging from 1 to 10. Error bars represent ±1 standard error from 500 bootstrap replicates. The optimal number of clusters is indicated by the value of k that maximizes the gap statistic

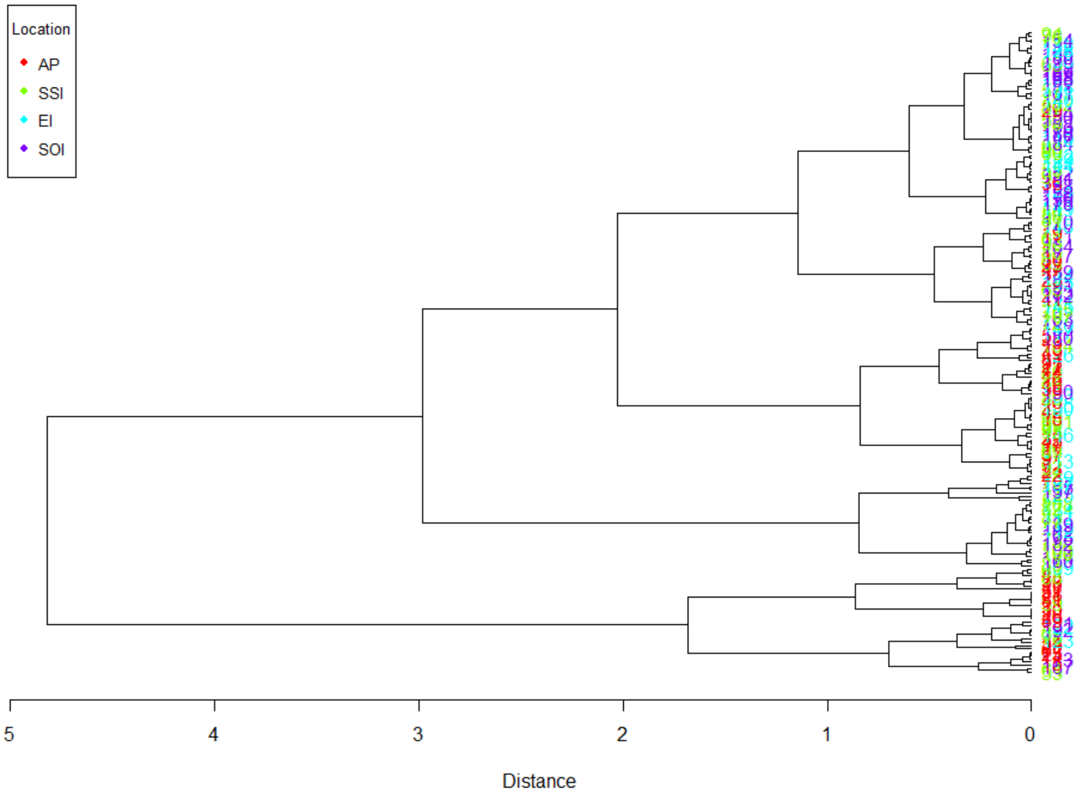


Fig S2. Hierarchical clustering dendrogram of chinstrap penguin (*Pygoscelis antarcticus*) colony sizes. The dendrogram was generated using complete linkage clustering based on Euclidean distances of log-transformed colony size data. Each leaf represents an individual colony, and the height of branch points indicates the distance at which clusters are merged.

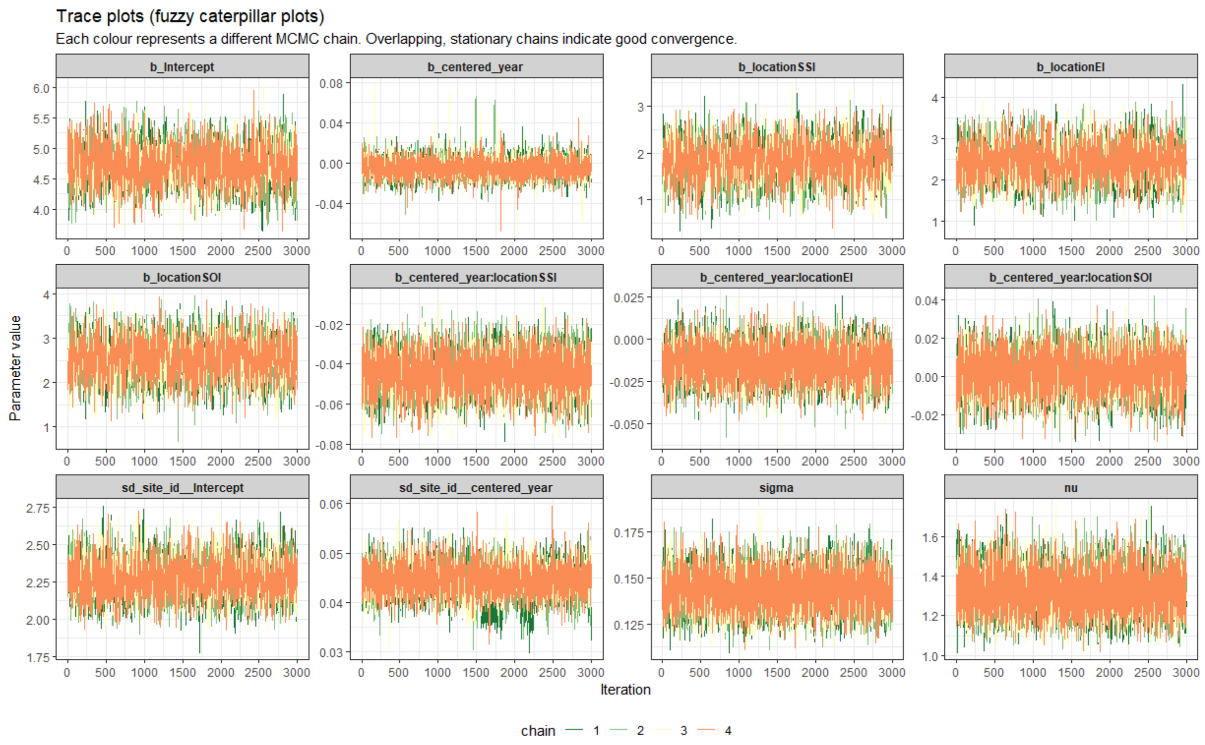


Fig S3 Convergence diagnostics for Bayesian Student-t interaction model. Trace plots of MCMC draws for all model parameters. Each panel displays four independent chains over 4000 iterations (1000 warmup). Visual inspection confirms adequate mixing and stationarity, supporting the reliability of posterior inferences. Parameter abbreviations: b_ = population-level effects, sd_ = standard deviations of random effects, sigma = residual standard deviation, nu = Student-t degrees of freedom.

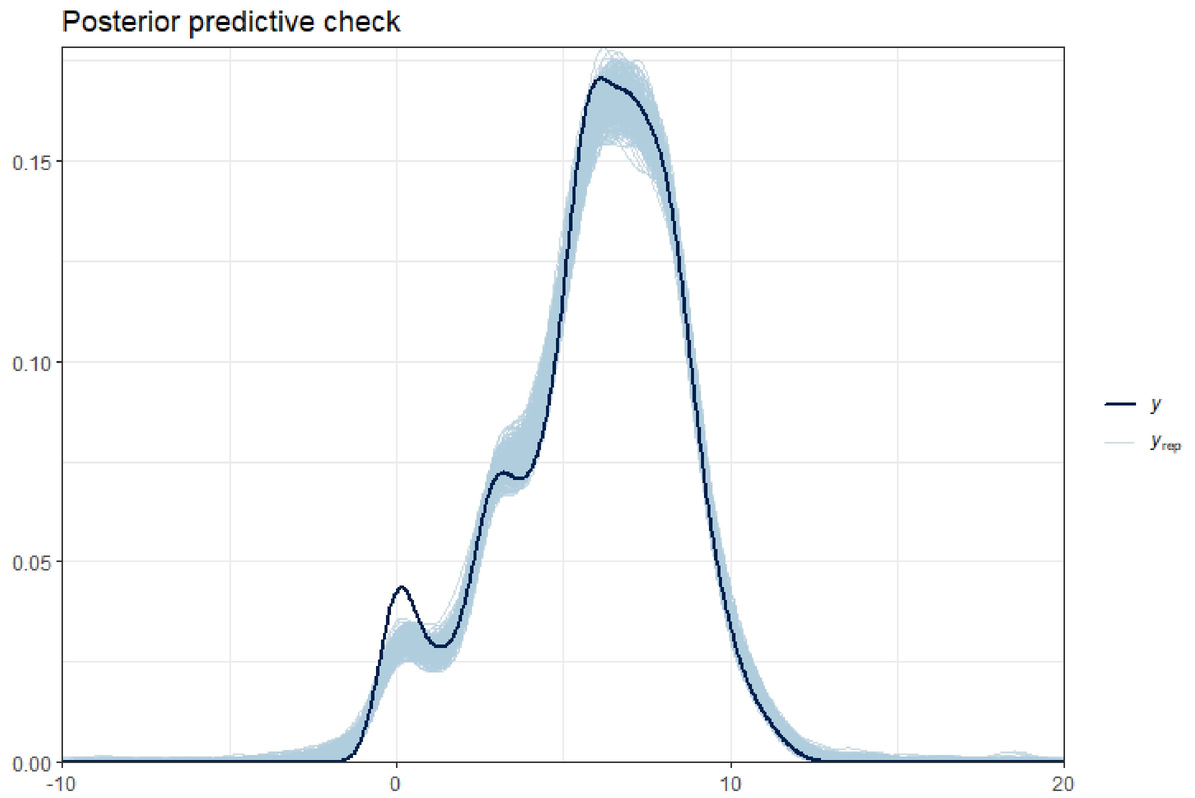


Fig S4. Posterior predictive check for Bayesian Student-t model. The plot compares the observed log-transformed penguin counts (dark blue line) with 500 simulated datasets drawn from the posterior predictive distribution (light blue lines). Good model fit is indicated by the observed data falling within the range of simulated datasets.

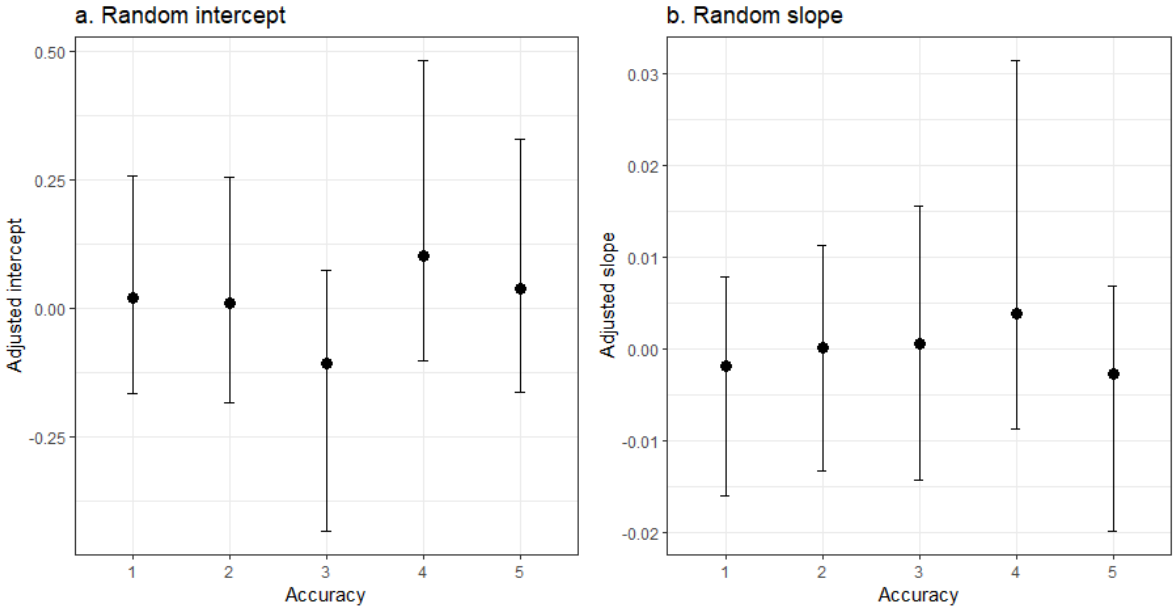


Fig S5. Accuracy-level random effects from the Bayesian Student-t interaction model. Panel (a) displays the random intercept adjustments for each accuracy level (1-5), representing accuracy-specific deviations from the global intercept. Panel (b) shows the random slope adjustments for each accuracy level, representing accuracy-specific deviations from the global effect of centered year. Points are posterior means, and error bars indicate 95% credible intervals
