## Supplementary File 2 for "A Bayesian hierarchical analysis of trends and risk to identify regional conservation priorities of Chinstrap Penguins": Supplemental_file_2_Methodological_Appendix.html


### Methodological Appendix

###### Lucas Krüger

###### Luis Adasme

###### Carlos Montenegro

###### Draft methodological appendix

**Companion to:** *A Bayesian hierarchical analysis
of trends and risk to identify regional conservation priorities of
Chinstrap Penguins*.

### 1 Purpose and scope

This appendix provides a formal and reproducible description of the
Bayesian hierarchical model used to estimate temporal trends in
Chinstrap Penguin nest counts across CCAMLR Subareas 48.1 and 48.2. It
describes the observation model, hierarchical linear predictor,
random-effects structure, prior distributions, posterior computation,
conservation-risk metrics, regional aggregation, compensation analysis,
and model diagnostics.

The principal inferential targets are:

1. annual trends on the logarithmic scale at global, location, and
   colony levels;
2. percentage change over three generations, defined as 28.2
   years;
3. posterior probabilities that declines exceed the 30% and 50% IUCN
   thresholds; and
4. the probability that increases in some colonies compensate for
   losses in declining colonies.

### 2 Statistical and mathematical nomenclature

This section defines the notation used throughout the appendix. A
**subscript** identifies a unit or observation, a
**superscript in parentheses** identifies a posterior draw,
and a Greek letter usually denotes an unknown model parameter.

| Symbol | Meaning |
| --- | --- |
| \(i\) | Colony index, \(i=1,\ldots,J\), with \(J=194\). |
| \(j\) | Observation index within colony, with a total of \(n=1{,}072\) records. |
| \(t\_{ij}\) | Calendar year of observation \(j\) from colony \(i\). |
| \(x\_{ij}\) | Centered calendar year used as the time covariate. |
| \(y\_{ij}\) | Observed number of nests. |
| \(z\_{ij}=\log(y\_{ij})\) | Natural logarithm of the observed nest count. |
| \(\ell(i)\) | Geographic location containing colony \(i\): AP, SSI, EI, or SOI. |
| \(a(i,j)\) | Accuracy class assigned to observation \((i,j)\), from 1 to 5. |
| \(\bar N\_i\) | Long-term mean nest count for colony \(i\). |
| \(\alpha\) | Population-level intercept. |
| \(\beta\) | Population-level annual slope for the reference location. |
| \(\gamma\_{\ell}\) | Location-specific intercept contrast. |
| \(\delta\_{\ell}\) | Location-specific slope contrast. |
| \(u\_i\) | Colony-specific deviation from the population intercept. |
| \(v\_i\) | Colony-specific deviation from the population slope. |
| \(w\_a\) | Effect associated with accuracy class \(a\). |
| \(\mu\_{ij}\) | Expected log nest count for observation \((i,j)\). |
| \(\sigma\) | Residual scale of the Student-\(t\) observation model. |
| \(\nu\) | Degrees of freedom of the Student-\(t\) distribution. |
| \(\sigma\_u,\sigma\_v\) | Standard deviations of colony intercept and slope effects. |
| \(\rho\) | Correlation between colony random intercepts and slopes. |
| \(\boldsymbol\Theta\) | Collective notation for all unknown parameters and random effects. |
| \(d\) | Posterior-draw index, \(d=1,\ldots,D\). |
| \(\mathbb{1}[\cdot]\) | Indicator function: 1 when the condition is true and 0 otherwise. |
| \(\propto\) | “Proportional to”; equality up to a normalizing constant. |
| \(\sim\) | “Is distributed as”. |
| \(\mid\) | “Conditional on” or “given”. |

The notation \(b^{(d)}\), for
example, means the value of parameter \(b\) in posterior draw \(d\). It is not an exponent.

### 3 Data structure and indexing

The analysis uses 1,072 nest-count observations from 194 colonies
collected between 1970 and 2024. Colonies are grouped into four
locations: Antarctic Peninsula (AP), South Shetland Islands (SSI),
Elephant Islands (EI), and South Orkney Islands (SOI). Each survey
record also belongs to one of five accuracy classes.

#### 3.1 Centering calendar year

Calendar year is centered at 1997, approximately the mean survey
year:

\[\begin{equation}
x\_{ij}=t\_{ij}-1997.
\tag{1}
\end{equation}\]

Centering does not change the estimated temporal slopes. It improves
numerical conditioning and makes the intercept interpretable: when \(x\_{ij}=0\), the observation year is
1997.

### 4 Observation model

Nest counts are positive and strongly right-skewed. The analysis
therefore models their natural logarithm:

\[
z\_{ij}=\log(y\_{ij}).
\]

Conditional on the expected log count and the residual parameters,
the observation model is

\[\begin{equation}
z\_{ij}\mid\mu\_{ij},\sigma,\nu
\sim \operatorname{Student\text{-}t}(\nu,\mu\_{ij},\sigma).
\tag{2}
\end{equation}\]

In this parameterization:

- \(\mu\_{ij}\) is the location
  parameter and represents the expected log count;
- \(\sigma>0\) is a scale
  parameter controlling residual dispersion; and
- \(\nu>0\) is the
  degrees-of-freedom parameter controlling tail thickness.

A Student-\(t\) distribution has
heavier tails than a Normal distribution when \(\nu\) is small and is therefore less
sensitive to unusually large or small observations. As \(\nu\) increases, it approaches a Gaussian
distribution.

#### 4.1 Likelihood contribution

Conditional on all model parameters and random effects, observations
are assumed independent. The likelihood is therefore

\[\begin{equation}
L(\boldsymbol\Theta;\mathbf z)
=
\prod\_{i=1}^{J}\prod\_{j=1}^{n\_i}
f\_t\!\left(z\_{ij}\mid\mu\_{ij},\sigma,\nu\right),
\tag{3}
\end{equation}\]

where \(f\_t(\cdot)\) denotes the
Student-\(t\) probability density and
\(n\_i\) is the number of observations
from colony \(i\).

The product symbol \(\prod\) means
that the individual density contributions are multiplied. Equivalently,
the log-likelihood is obtained by summing their logarithms. Conditional
independence means that any residual temporal or spatial dependence not
explained by location, colony, year, or survey accuracy is not modeled
explicitly.

### 5 Hierarchical linear predictor

The expected log count is modeled as

\[\begin{equation}
\mu\_{ij}
=
\alpha+\beta x\_{ij}
+\gamma\_{\ell(i)}
+\delta\_{\ell(i)}x\_{ij}
+u\_i+v\_i x\_{ij}
+w\_{a(i,j)}.
\tag{4}
\end{equation}\]

Each component has a distinct interpretation:

| Term | Interpretation |
| --- | --- |
| \(\alpha\) | Expected log nest count in the reference location, AP, in 1997. |
| \(\beta\) | Annual log-scale trend in AP. |
| \(\gamma\_{\ell(i)}\) | Difference in intercept between location \(\ell(i)\) and AP. |
| \(\delta\_{\ell(i)}\) | Difference in annual trend between location \(\ell(i)\) and AP. |
| \(u\_i\) | Colony-specific intercept deviation. |
| \(v\_i\) | Colony-specific slope deviation. |
| \(w\_{a(i,j)}\) | Systematic effect associated with the observation’s accuracy class. |

The fixed effects \(\alpha\), \(\beta\), \(\gamma\_{\ell}\), and \(\delta\_{\ell}\) describe population-level
patterns. The random effects \(u\_i\),
\(v\_i\), and \(w\_a\) describe group-specific departures
from those patterns.

#### 5.1 Reference-category coding and location-specific slopes

AP is the reference location. Therefore its annual trend is

\[\begin{equation}
\beta\_{\mathrm{AP}}=\beta,
\qquad
\beta\_{\ell}=\beta+\delta\_{\ell},
\quad \ell\in\{\mathrm{SSI},\mathrm{EI},\mathrm{SOI}\}.
\tag{5}
\end{equation}\]

The parameter \(\delta\_{\ell}\) is
not the complete trend for location \(\ell\); it is the difference between that
location’s trend and the AP trend.

#### 5.2 Colony-specific trends and partial pooling

The annual trend for colony \(i\)
is

\[\begin{equation}
\beta\_i=\beta+\delta\_{\ell(i)}+v\_i.
\tag{6}
\end{equation}\]

Thus, a colony trend is formed by adding the AP reference slope, the
appropriate location contrast, and the colony-specific random-slope
deviation.

The hierarchical structure produces **partial pooling**.
Colonies with abundant information may depart substantially from the
corresponding location trend when the data support that departure.
Colonies with only a few observations are shrunk more strongly toward
the location-level trend. This shrinkage is what is meant by “borrowing
strength” across colonies.

### 6 Random-effects distributions

#### 6.1 Colony intercepts and slopes

The colony-specific intercept and slope are modeled jointly:

\[\begin{equation}
\begin{pmatrix}u\_i\\v\_i\end{pmatrix}
\sim
\operatorname{MVN}\_2\!\left[
\begin{pmatrix}0\\0\end{pmatrix},
\boldsymbol\Sigma\_{\mathrm{site}}
\right].
\tag{7}
\end{equation}\]

Here \(\operatorname{MVN}\_2\)
denotes a two-dimensional multivariate Normal distribution. Its
covariance matrix is decomposed as

\[\begin{equation}
\boldsymbol\Sigma\_{\mathrm{site}}
=
\mathbf D\mathbf R\mathbf D,
\qquad
\mathbf D=\operatorname{diag}(\sigma\_u,\sigma\_v),
\tag{8}
\end{equation}\]

with correlation matrix

\[\begin{equation}
\mathbf R=
\begin{pmatrix}
1 & \rho\\
\rho & 1
\end{pmatrix}.
\tag{9}
\end{equation}\]

The decomposition separates marginal standard deviations from
correlation:

- \(\sigma\_u\) measures
  between-colony variability in intercepts;
- \(\sigma\_v\) measures
  between-colony variability in annual trends; and
- \(\rho\) measures association
  between colony intercept and slope deviations.

A positive \(\rho\) means that
colonies with higher-than-expected baseline abundance tend to have more
positive, or less negative, trends after accounting for location. A
negative \(\rho\) indicates the
opposite.

#### 6.2 Accuracy effects

The accuracy-class effects are modeled as

\[\begin{equation}
w\_a\sim\operatorname{Normal}(0,\sigma\_{\mathrm{acc}}^2),
\qquad a=1,\ldots,5.
\tag{10}
\end{equation}\]

This is a random-intercept formulation. It allows the average
observed count to differ systematically among survey-quality classes
after controlling for colony, location, and year. It does
**not** assign a known observation-error variance to each
class.

If the fitted model includes an accuracy-by-year random slope,
equation (4) must also contain a term such as \(s\_a x\_{ij}\), and the joint distribution of
\(w\_a\) and \(s\_a\) must be specified.

### 7 Prior distributions

Weakly informative priors regularize the model while allowing the
data to dominate when information is strong.

| Parameter | Prior | Interpretation |
| --- | --- | --- |
| \(\alpha\) | \(\operatorname{Normal}(0,2)\) | Prior for the reference intercept on the log scale. |
| \(\beta,\gamma,\delta\) | \(\operatorname{Normal}(0,1)\) | Priors for slopes and location contrasts. |
| \(\sigma\_u,\sigma\_v\) | \(\operatorname{Exponential}(2)\) | Positive priors shrinking hierarchical standard deviations toward zero. |
| \(\sigma\_{\mathrm{acc}}\) | \(\operatorname{Student\text{-}t}^{+}(3,0,1)\) | Half-Student-\(t\) prior for the accuracy-effect standard deviation. |
| \(\mathbf R\) | \(\operatorname{LKJ}(4)\) | Prior on the random-effects correlation matrix. |
| \(\nu\) | \(\operatorname{Gamma}(2,0.1)\) | Prior for Student-\(t\) degrees of freedom, using shape-rate parameterization. |
| \(\sigma\) | \(\operatorname{Student\text{-}t}^{+}(3,0,1)\) | Half-Student-\(t\) prior for residual scale. |

#### 7.1 Explanation of the distribution names

**Normal distribution.** A symmetric continuous
distribution described by a mean and standard deviation. A prior \(\operatorname{Normal}(0,1)\) expresses that
values near zero are more plausible a priori, while positive and
negative departures remain possible.

**Exponential distribution.** A positive continuous
distribution. The notation \(\operatorname{Exponential}(2)\) uses rate
2, giving prior mean \(1/2\). It places
greatest density near zero but retains a right tail for larger standard
deviations.

**Half-Student-\(t\)
distribution.** The superscript \(+\) means that only positive values are
allowed. This is appropriate for scale and standard-deviation
parameters, which cannot be negative. Heavy tails permit occasional
large values without placing excessive prior mass on them.

**Gamma distribution.** A positive continuous
distribution. Under the shape-rate convention, \(\nu\sim\operatorname{Gamma}(2,0.1)\) has
prior mean \(2/0.1=20\). Because \(\nu\) controls tail thickness, this prior
allows both robust heavy-tailed models and models close to Gaussian.

**LKJ distribution.** The LKJ distribution is a prior
over valid correlation matrices, not over a single regression
coefficient. In the two-dimensional case it induces a prior on \(\rho\). With concentration parameter 4, it
favors correlations near zero while still allowing positive or negative
correlations supported by the data. Larger LKJ concentration values
imply stronger regularization toward the identity matrix.

#### 7.2 Interpretation on the original count scale

Because the response is modeled on the natural-log scale, an annual
coefficient \(b\) is additive on the
log scale but multiplicative on the original count scale. The
corresponding annual percentage change is

\[\begin{equation}
100\left[\exp(b)-1\right].
\tag{11}
\end{equation}\]

For example, \(b=-0.02\) implies a
multiplicative factor \(\exp(-0.02)\approx0.9802\), or an annual
decline of approximately 1.98%.

#### 7.3 Prior predictive checking

Priors should be evaluated through the data patterns they imply.
Prior predictive simulation should therefore be used to verify that the
model generates ecologically plausible colony sizes and temporal
trajectories before conditioning on observed data.

This is especially important for annual trends. A \(\operatorname{Normal}(0,1)\) prior on an
unscaled yearly slope permits extremely large year-to-year changes on
the original abundance scale. Its practical informativeness must
therefore be assessed using the actual units and range of \(x\_{ij}\).

### 8 Posterior distribution

Let \(\boldsymbol\Theta\) denote all
fixed effects, random effects, standard deviations, correlations,
residual scale, and degrees of freedom. Bayes’ theorem gives

\[\begin{equation}
p(\boldsymbol\Theta\mid\mathbf z)
=
\frac{p(\mathbf z\mid\boldsymbol\Theta)p(\boldsymbol\Theta)}
{\int p(\mathbf
z\mid\boldsymbol\Theta)p(\boldsymbol\Theta)\,d\boldsymbol\Theta}.
\tag{12}
\end{equation}\]

The numerator combines the likelihood and prior. The denominator is
the **marginal likelihood**, which normalizes the posterior
so that it integrates to one. For posterior sampling with HMC, it is
sufficient to know the posterior up to proportionality:

\[\begin{equation}
\begin{split}
p(\boldsymbol\Theta\mid\mathbf z)
\propto {}& L(\boldsymbol\Theta;\mathbf z)
\,p(\alpha)\,p(\beta,\boldsymbol\gamma,\boldsymbol\delta)
\,p(\mathbf u,\mathbf v\mid\boldsymbol\Sigma\_{\mathrm{site}})\\
&\times p(\mathbf w\mid\sigma\_{\mathrm{acc}})
\,p(\boldsymbol\Sigma\_{\mathrm{site}})
\,p(\sigma)\,p(\nu).
\end{split}
\tag{13}
\end{equation}\]

### 9 Posterior computation with HMC and NUTS

The model was fitted using `brms`, which generates Stan
code and samples from the joint posterior using Hamiltonian Monte Carlo
(HMC) with the No-U-Turn Sampler (NUTS).

HMC augments the parameter vector \(\boldsymbol\Theta\) with an auxiliary
momentum vector \(\mathbf p\) and
defines the Hamiltonian

\[\begin{equation}
H(\boldsymbol\Theta,\mathbf p)
=U(\boldsymbol\Theta)+K(\mathbf p),
\tag{14}
\end{equation}\]

where

\[\begin{equation}
U(\boldsymbol\Theta)=-\log p(\boldsymbol\Theta\mid\mathbf z),
\qquad
K(\mathbf p)=\frac{1}{2}\mathbf p^{\mathsf T}\mathbf M^{-1}\mathbf p.
\tag{15}
\end{equation}\]

\(U\) is the potential energy, equal
to the negative log posterior. \(K\) is
the kinetic energy, and \(\mathbf M\)
is the mass matrix. HMC uses gradients of the log posterior to construct
distant proposals with high acceptance probability, reducing random-walk
behavior. NUTS automatically chooses the trajectory length and stops
when the simulated path begins to turn back toward its starting
point.

#### 9.1 Chains, warmup, and retained draws

Four independent chains were run for 4,000 iterations each. The first
1,000 iterations per chain were used for warmup, during which Stan
adapted the step size and mass matrix. The retained sample contained

\[\begin{equation}
4\times(4{,}000-1{,}000)=12{,}000
\quad\text{posterior draws}.
\tag{16}
\end{equation}\]

Each retained draw is one joint realization of all parameters.
Derived quantities must therefore be calculated draw by draw so that
posterior dependence is retained and uncertainty is propagated
correctly.

### 10 From annual trend to three-generation change

For a log-linear annual trend \(b\),
expected abundance is proportional to \(\exp(bt)\). Over an interval of \(G=28.2\) years, the abundance ratio is
\(\exp(Gb)\). The percentage change
over three generations is therefore

\[\begin{equation}
\Delta\_G(b)=100\left[\exp(Gb)-1\right],
\qquad G=28.2.
\tag{17}
\end{equation}\]

For posterior draw \(d\),

\[\begin{equation}
\Delta\_G^{(d)}
=100\left[\exp\left(28.2\,b^{(d)}\right)-1\right].
\tag{18}
\end{equation}\]

Posterior medians and credible intervals should be calculated from
the transformed draws \(\Delta\_G^{(d)}\). Because the
transformation is nonlinear, transforming a summary of \(b\) is not generally equivalent to
summarizing the transformed draws.

#### 10.1 Interpretation and projection assumption

This quantity represents the change implied by maintaining the
estimated average annual trend for 28.2 years. It is a standardized
projection for IUCN assessment, not a mechanistic forecast. It does not
account for future changes in climate, prey availability, fishing
pressure, density dependence, or nonlinear colony dynamics.

### 11 Posterior probabilities of IUCN decline thresholds

The probability of a decline of at least 30% is estimated directly
from posterior draws:

\[\begin{equation}
P\_{30}
=P(\Delta\_G\leq-30\mid\mathbf z)
\approx
\frac{1}{D}\sum\_{d=1}^{D}
\mathbb{1}\!\left[\Delta\_G^{(d)}\leq-30\right].
\tag{19}
\end{equation}\]

Similarly,

\[\begin{equation}
P\_{50}
=P(\Delta\_G\leq-50\mid\mathbf z)
\approx
\frac{1}{D}\sum\_{d=1}^{D}
\mathbb{1}\!\left[\Delta\_G^{(d)}\leq-50\right].
\tag{20}
\end{equation}\]

The indicator function \(\mathbb{1}[A]\) equals 1 when statement
\(A\) is true and 0 otherwise. Thus,
these estimates are simply the proportions of posterior draws crossing
the respective thresholds. For example, \(P\_{30}=0.914\) means that 91.4% of
posterior draws imply a decline of at least 30% over 28.2 years,
conditional on the model, data, and priors.

### 12 Population-weighted regional aggregation

An unweighted mean would give a small colony the same influence as a
colony containing tens of thousands of nests. Population weights are
therefore defined using long-term mean colony counts:

\[\begin{equation}
w\_i=\frac{\bar N\_i}{\sum\_{k=1}^{J}\bar N\_k},
\qquad
\sum\_{i=1}^{J}w\_i=1.
\tag{21}
\end{equation}\]

#### 12.1 Log-scale weighted aggregation

For posterior draw \(d\), the
manuscript defines the population-weighted regional log trend as

\[\begin{equation}
\beta\_{\mathrm{pop}}^{(d)}
=\sum\_{i=1}^{J}w\_i\beta\_i^{(d)}.
\tag{22}
\end{equation}\]

The corresponding three-generation percentage change is

\[\begin{equation}
\Delta\_{\mathrm{pop}}^{(d)}
=100\left[
\exp\left(28.2\,\beta\_{\mathrm{pop}}^{(d)}\right)-1
\right].
\tag{23}
\end{equation}\]

This procedure averages log growth rates before exponentiation and
therefore produces a weighted geometric aggregation.

#### 12.2 Direct abundance-scale aggregation

A more direct population-scale calculation projects each colony
separately and then sums projected abundance:

\[\begin{equation}
\Delta\_{\mathrm{pop,abund}}^{(d)}
=100\left\{
\frac{\sum\_{i=1}^{J}\bar N\_i
\exp\left(28.2\,\beta\_i^{(d)}\right)}
{\sum\_{i=1}^{J}\bar N\_i}
-1
\right\}.
\tag{24}
\end{equation}\]

Equations (23) and (24) are similar when colony trends are
homogeneous, but they may differ substantially under strong spatial
heterogeneity. The authors should verify which formulation was
implemented and use the same definition in the manuscript, appendix,
tables, figures, and code.

### 13 Compensation analysis

For each posterior draw, the projected absolute change in colony
\(i\) is

\[\begin{equation}
A\_i^{(d)}
=\bar N\_i\left[
\exp\left(28.2\,\beta\_i^{(d)}\right)-1
\right].
\tag{25}
\end{equation}\]

Positive and negative contributions are then separated:

\[\begin{equation}
\operatorname{Increase}^{(d)}
=\sum\_{i:\,\beta\_i^{(d)}>0}A\_i^{(d)},
\tag{26}
\end{equation}\]

\[\begin{equation}
\operatorname{Loss}^{(d)}
=\sum\_{i:\,\beta\_i^{(d)}<0}\left|A\_i^{(d)}\right|,
\tag{27}
\end{equation}\]

and

\[\begin{equation}
\operatorname{NetChange}^{(d)}
=\operatorname{Increase}^{(d)}-\operatorname{Loss}^{(d)}
=\sum\_{i=1}^{J}A\_i^{(d)}.
\tag{28}
\end{equation}\]

The compensation probability is

\[\begin{equation}
P\_{\mathrm{comp}}
=P(\operatorname{NetChange}>0\mid\mathbf z)
\approx
\frac{1}{D}\sum\_{d=1}^{D}
\mathbb{1}\!\left[\operatorname{NetChange}^{(d)}>0\right].
\tag{29}
\end{equation}\]

All colony quantities within a draw must be computed from the same
joint posterior draw. This preserves posterior correlations among colony
trends.

### 14 MCMC convergence and sampling diagnostics

#### 14.1 Rank-normalized split \(\widehat R\)

The statistic \(\widehat R\)
compares within-chain and between-chain variability. Values close to 1
indicate that independently initialized chains are exploring the same
posterior region. The analysis required \(\widehat R\leq1.01\) for all monitored
quantities.

A satisfactory \(\widehat R\)
supports convergence of the sampler but does not establish that the
scientific model is correctly specified.

#### 14.2 Effective sample size

MCMC draws are autocorrelated, so 12,000 stored draws contain less
information than 12,000 independent observations. The effective sample
size (ESS) estimates the equivalent number of independent draws.

- **Bulk ESS** assesses estimation of the center of the
  posterior.
- **Tail ESS** assesses estimation of posterior quantiles
  and tail probabilities.

Large values support stable estimates of medians, credible intervals,
and threshold probabilities.

#### 14.3 Divergent transitions

A divergent transition indicates that the numerical HMC trajectory
could not accurately follow a region of high posterior curvature.
Divergences may reveal funnel-shaped geometry, weak identification, poor
scaling, or an unsuitable parameterization.

The manuscript reports one divergence, approximately 0.01% of
retained draws. Its location should be inspected with pair plots.
Possible responses include increasing `adapt_delta`, using a
non-centered parameterization, strengthening priors, or rescaling
predictors.

### 15 Posterior predictive assessment

For each posterior draw, a replicated dataset is generated from the
fitted observation model:

\[\begin{equation}
z\_{\mathrm{rep}}^{(d)}
\sim p\!\left(z\_{\mathrm{rep}}\mid\boldsymbol\Theta^{(d)}\right).
\tag{30}
\end{equation}\]

The replicated data should be compared with the observed data.
Density overlays are useful but insufficient on their own. Recommended
checks include:

- distributions and extreme values of counts;
- colony-specific residual patterns;
- temporal autocorrelation;
- trends by location; and
- distributions of within-colony changes.

An aggregated density may fit well even when the model fails to
represent important temporal or colony-level structure.

### 16 Leave-one-out cross-validation and Pareto-\(k\)

Pareto-smoothed importance-sampling leave-one-out cross-validation
(PSIS-LOO) approximates predictive performance for observations omitted
one at a time without fitting the model \(n\) separate times. The Pareto shape
parameter \(k\) diagnoses the stability
of the importance-sampling approximation.

| Pareto-\(k\) range | Typical interpretation |
| --- | --- |
| \(k\leq0.5\) | Reliable approximation. |
| \(0.5<k\leq0.7\) | Usually usable, but observations are increasingly influential. |
| \(0.7<k\leq1\) | Potentially unreliable; moment matching or exact refitting is advisable. |
| \(k>1\) | Importance-sampling estimate is unreliable. |

The manuscript reports 26 observations with \(k>0.7\) and states that moment matching
was used. The influential records should be examined to determine
whether they represent data errors, legitimate but rare colony
trajectories, or inadequacy of the assumed model.

### 17 Bayesian \(R^2\)

For each posterior draw, Bayesian \(R^2\) can be written as

\[\begin{equation}
R^{2(d)}
=
\frac{\operatorname{Var}\!\left(\boldsymbol\mu^{(d)}\right)}
{\operatorname{Var}\!\left(\boldsymbol\mu^{(d)}\right)
+\operatorname{Var}\!\left(\mathbf z-\boldsymbol\mu^{(d)}\right)}.
\tag{31}
\end{equation}\]

The numerator measures variation in fitted values, and the
denominator adds residual variation. A reported value near 0.946
indicates strong in-sample explanatory performance on the log-count
scale. It does not establish causality, ensure accurate extrapolation,
or rule out misspecification.
