## Supplementary File 3 for "A Bayesian hierarchical analysis of trends and risk to identify regional conservation priorities of Chinstrap Penguins": Supplementary_File_3_Rcode.R

```
#  Load libraries -------------------------------------------------------
library(ggplot2)
library(dplyr)
library(tidyr)
library(brms)
library(bayesplot)
library(tidybayes)
library(patchwork)
library(stringr)
library(purrr)
library(cluster)
library(factoextra)
library(ggplotify)
library(dendextend)

gc()

# -----Read and prepare data ---------------------------------------------------
all <- read.csv("AllCounts_V_4_4.csv") %>% 
  filter(season_starting >= 1970)%>%
  select(site_name,site_id,cammlr_region,lat=latitude_epsg_4326,
         lon=longitude_epsg_4326,season_starting,penguin_count,
         accuracy,count_type,common_name)


GENERATION_LENGTH <- 9.4   # chinstrap penguin generation length (years)

# Initial data preparation (including zeros, accuracy <=5)
model_data <- all %>%
  filter(common_name == "chinstrap penguin",
         count_type == "nests",
         accuracy <= 5) %>%        # keep all accuracy levels, zeros included
                                  # 
  group_by(site_id) %>%
  mutate(
    mean_site_size = mean(penguin_count, na.rm = TRUE),
    centered_year = season_starting - mean(season_starting),
    log_count = log(penguin_count + 1),   # log(0+1)=0
    site_id = as.factor(site_id),
    accuracy_factor = as.factor(accuracy)
  ) %>%
  group_by(site_id) %>%
  filter(n_distinct(season_starting) >= 2) %>%   # at least 2 years per site
  
  ungroup()


 
 # a simple measure of change to identify sites to inspect data


inspect_colonies <- model_data %>%
  group_by(site_id) %>%
  summarise(
    first_year = min(season_starting),
    last_year = max(season_starting),
    first_count = penguin_count[which.min(season_starting)],
    last_count = penguin_count[which.max(season_starting)],
    max_count = max(penguin_count),
    min_count = min(penguin_count),
    n_obs = n(),
    accuracy_first = accuracy[which.min(season_starting)],
    accuracy_last = accuracy[which.max(season_starting)],
    
    .groups = "drop"
  ) %>%
  mutate(
    interval = last_year - first_year,
    total_perc_change = ((last_count / first_count) - 1) * 100,
    annual_perc_change = total_perc_change / interval,
    nest_change = last_count - first_count,
    annual_nest_change = nest_change / interval,
    # Flag unreasonable changes
    flag = case_when(
      interval == 0 ~ "Same year only",
      n_obs < 3 ~ "Few observations",
      abs(annual_perc_change) > 20 ~ "Extreme annual change (>20% per year)",
      annual_nest_change > 50000 ~ "Extreme annual nest change (>50,000 nests/year)",
      accuracy_first >= 4 | accuracy_last >= 4 ~ "Low accuracy",
      TRUE ~ "OK"
    )
  ) %>%
  arrange(desc(abs(annual_perc_change)))

# View suspicious colonies
inspect_colonies %>%
  filter(flag != "OK") %>%
  select(site_id, first_year, last_year, first_count, last_count, 
         annual_perc_change, annual_nest_change, flag) %>%
  print(n = 50)


ggplot(inspect_colonies,aes(reorder(site_id,annual_nest_change),annual_nest_change))+geom_point()+
  coord_flip()


lower_end<-inspect_colonies%>%
  filter(annual_nest_change<(-500))


upper_end<-inspect_colonies%>%
  filter(annual_nest_change>(200))


model_data%>%
  filter(site_id %in% c(unique(lower_end$site_id)))%>%
  ggplot(aes(season_starting,penguin_count,colour=accuracy_factor))+geom_point()+
  facet_wrap(site_id~.)

# seems these are fine


model_data%>%
  filter(site_id %in% c(unique(upper_end$site_id)))%>%
  ggplot(aes(season_starting,penguin_count,colour=accuracy_factor))+geom_point()+
  facet_wrap(site_id~.)

# Harmony Point have had issues in past estimations, so let's check

model_data%>%
  filter(site_id %in% c("HARM"))%>%
  ggplot(aes(season_starting,penguin_count,colour=accuracy_factor))+geom_point()+
  facet_wrap(site_id~.)

# Remove single observation: Harmony Point 1971
model_data_clean <- model_data %>%
  filter(!(site_id == "HARM" & season_starting == 1971))
# 
message("Original observations: ", nrow(model_data))
message("Cleaned observations: ", nrow(model_data_clean))


table(model_data_clean$cammlr_region)


# classify sites based on regions


model_data_clean$location[model_data_clean$cammlr_region=="48.2"]<-"SOI"

model_data_clean$location[model_data_clean$lon<(-50) &
                            model_data_clean$lon>(-56.75) ]<-"EI"

model_data_clean$location[is.na(model_data_clean$location) &
                            model_data_clean$lat>(-63.3) ]<-"SSI"


model_data_clean$location[is.na(model_data_clean$location) ]<-"AP"

table(model_data_clean$location)

model_data_clean$location<-factor(model_data_clean$location,
                              levels=c("AP","SSI","EI","SOI"))


# Hierarchical clustering to define size categories -------------------

colony_cluster_data <- model_data_clean %>%
  group_by(location,site_id) %>%
  summarise(mean_size = mean(penguin_count, na.rm = TRUE)) %>%
  mutate(log_size = log10(mean_size + 1),
         scaled_size = scale(log_size))


# Hierarchical clustering with complete linkage


# Compute the gap statistic for up to 10 possible clusters
set.seed(123)
gap_stat <- clusGap((colony_cluster_data$scaled_size),
                    FUN = kmeans,
                    nstart = 25,
                    K.max = 10,
                    B = 500)

# Plot the gap statistic
plot(gap_stat,type = "b", xlab = "k", ylab = expression(Gap[k]),
     main = "", do.arrows = TRUE,ylim = c(0, 0.4),
     arrowArgs = list(col="red3", length=1/16, angle=90, code=3))

hc <- hclust(dist(colony_cluster_data$log_size), method = "complete")


# First, create a color vector based on location
locations <- colony_cluster_data$location
unique_locations <- unique(locations)
location_colors <- setNames(rainbow(length(unique_locations)), unique_locations)
leaf_colors <- location_colors[locations]

# color the dendrogram leaves by location
dend <- as.dendrogram(hc)
labels_colors(dend) <- leaf_colors[order.dendrogram(dend)]

# plot with colored leaves
plot(dend, 
     horiz = TRUE, 
     main = "",
     xlab = "Distance",
     ylab = "",
     #leaflab = "textlike",  # Show labels
     cex = 0.1)  # Adjust text size

# add legend
legend("topleft", 
       legend = unique_locations,
       col = location_colors[unique_locations],
       pch = 16,
       title = "Location",
       cex = 0.8)

k_clusters <- 4

colony_cluster_data$size_cluster <- cutree(hc, k = k_clusters)

colony_cluster_data%>%
  group_by(size_cluster)%>%
  summarise(min=min(mean_size),
            max=max(mean_size))

# Map 7 clusters to 4 meaningful categories
colony_cluster_data$size_category <- case_when(
  # small clusters (clusters 2,3) → "small"
  colony_cluster_data$size_cluster %in% c(2,3) ~ "small",
  
  # Medium clusters (clusters 1,2) → "medium" 
  colony_cluster_data$size_cluster %in% c(1) ~ "medium",
  
  # Large cluster (cluster 6) → "large"
  colony_cluster_data$size_cluster == 4 ~ "large",
  

  TRUE ~ NA_character_
)

table(colony_cluster_data$size_category)

colony_cluster_data%>%
  group_by(size_category)%>%
  summarise(min=min(mean_size),
            max=max(mean_size))


ggplot(colony_cluster_data,aes(reorder(size_category,-mean_size),mean_size))+
  geom_boxplot()+
  scale_y_log10(labels=scales::comma_format())+
  theme_bw()+
  xlab("Colony size category")+
  ylab("Mean number of active nests")

##ggsave("colony_size_categories.pdf", width = 6, height = 4, dpi = 600)

colony_size_ranges<-colony_cluster_data%>%
  group_by(size_category)%>%
  summarise(min=min(mean_size),
            max=max(mean_size))

print(colony_size_ranges)

table(colony_cluster_data$size_category)


model_data_clean <- model_data_clean %>%
  left_join(colony_cluster_data[, c("site_id", "size_category")], by = "site_id")


table(model_data_clean$size_category)


# Ensure factor order
model_data_clean$size_category <- factor(model_data_clean$size_category,
                                         levels = c("small", "medium", "large"))


# Calculate the counts, this is going to be used at the end


count_data <- model_data_clean %>%
  group_by(location, size_category) %>%
  summarise(
    counts = n(),  # number of penguin count observations
    colonies = n_distinct(site_id),  # number of distinct colonies (adjust column name as needed)
    .groups = 'drop'
  )


# with the exception of AP that has no large colonies, all other sectors have 
# the three categories
# we can also see that the large colonies have the least amount of counts
 

# Bayesian model with Student-t distribution --------------------------

interaction_model_t <- brm(
  bf(log_count ~ centered_year * location +
       (centered_year | site_id) +
       (centered_year | accuracy_factor)),
  data = model_data_clean,
  family = student(),
  prior = c(
    prior(normal(0, 2), class = "Intercept"),
    prior(normal(0, 1), class = "b"),
    prior(exponential(2), class = "sd", group = "site_id"),
    prior(student_t(3, 0, 1), class = "sd", group = "accuracy_factor"),
    prior(lkj(4), class = "cor"),
    prior(gamma(2, 0.1), class = "nu")   # degrees of freedom
  ),
  chains = 4,
  iter = 4000,
  warmup = 1000,
  cores = 4,
  control = list(adapt_delta = 0.99, max_treedepth = 15),
  save_pars = save_pars(all = TRUE)
)

# Save model object 
saveRDS(interaction_model_t, "Results/interaction_model_t.rds")

# Model diagnostics ---------------------------------------------------
summary(interaction_model_t)
bayes_R2(interaction_model_t)


# 
# TRACE PLOTS (fuzzy caterpillar plots) for convergence check
# 
# Extract posterior draws as a matrix
posterior_draws <- as_draws_matrix(interaction_model_t)
parameters(interaction_model_t)


# Choose the most important parameters to monitor
trace_params <- c(
  "b_Intercept"  ,                                
  "b_centered_year" ,        
    
  "b_locationSSI"    ,                            
  "b_locationEI"     ,                            
  "b_locationSOI"     ,                           

  
  "b_centered_year:locationSSI" ,                 
  "b_centered_year:locationEI"  ,                 
  "b_centered_year:locationSOI" ,                 


      
  "sd_site_id__Intercept"   ,                     
  "sd_site_id__centered_year"  ,                  
  "sigma"  ,                                      
  "nu"  # degrees of freedom of the Student-t
)

# Create the trace plot
trace_plot <- mcmc_trace(posterior_draws,
                         pars = trace_params,
                         facet_args = list(ncol = 4, scales = "free")) +
  scale_color_manual(values = c("#1b7837", "#7fbf7b", "#ffffbf", "#fc8d59")) +
  labs(title = "Trace plots (fuzzy caterpillar plots)",
       subtitle = "Each colour represents a different MCMC chain. Overlapping, stationary chains indicate good convergence.",
       x = "Iteration",
       y = "Parameter value") +
  theme_bw() +
  theme(legend.position = "bottom",
        strip.background = element_rect(fill = "lightgray"),
        strip.text = element_text(face = "bold"))

# Display the plot
print(trace_plot)

# Save as PDF (high resolution)


# Posterior predictive check
y_rep <- posterior_predict(interaction_model_t, ndraws = 500)
ppc_dens_overlay(y = interaction_model_t$data$log_count, yrep = y_rep) +
  labs(title = "Posterior predictive check") +
  theme_bw()+xlim(-10,20)

# Save as PDF (high resolution)


# LOO with moment matching (requires save_pars = TRUE)
loo_result <- loo(interaction_model_t, moment_match = TRUE)
print(loo_result)

# Perform 10-fold cross-validation
# This will take longer but is more robust for problematic observations
# Increase the allowed size for parallel processing
options(future.globals.maxSize = 800 * 1024^2)  # 800 MB

# Now run kfold again
kfold_result <- kfold(interaction_model_t, K = 10, chains = 2, iter = 1000, warmup = 500)

# View results
print(kfold_result)


# loo_result <- loo(interaction_model_t, moment_match = TRUE)
# loo_compare(kfold_result, loo_result)


# Extract accuracy effects --------------------------------------------
accuracy_intercept <- interaction_model_t %>%
  spread_draws(r_accuracy_factor[accuracy, term]) %>%
  filter(term == "Intercept") %>%
  group_by(accuracy) %>%
  summarise(
    intercept_adj = mean(r_accuracy_factor),
    intercept_lower = quantile(r_accuracy_factor, 0.025),
    intercept_upper = quantile(r_accuracy_factor, 0.975)
  ) %>%
  mutate(accuracy = factor(accuracy, levels = 1:5))

accuracy_slope <- interaction_model_t %>%
  spread_draws(r_accuracy_factor[accuracy, term]) %>%
  filter(term == "centered_year") %>%
  group_by(accuracy) %>%
  summarise(
    slope_adj = mean(r_accuracy_factor),
    slope_lower = quantile(r_accuracy_factor, 0.025),
    slope_upper = quantile(r_accuracy_factor, 0.975)
  ) %>%
  mutate(accuracy = factor(accuracy, levels = 1:5))

accuracy_effects <- accuracy_intercept %>%
  left_join(accuracy_slope, by = "accuracy")

print(accuracy_effects)


ggplot(accuracy_effects,aes(as.factor(accuracy),intercept_adj))+
  geom_errorbar(aes(ymin=intercept_lower,ymax=intercept_upper),width=0.1)+
  geom_point(size=3)+
  theme_bw()+
  xlab("Accuracy")+ ylab("Adjusted intercept")+
  ggtitle(label="a. Random intercept")+

ggplot(accuracy_effects,aes(as.factor(accuracy),slope_adj))+
  geom_errorbar(aes(ymin=slope_lower,ymax=slope_upper),width=0.1)+
  geom_point(size=3)+
  theme_bw()+
  xlab("Accuracy")+ ylab("Adjusted slope")+
  ggtitle(label="b. Random slope")


#ggsave("Results/accuracy_effects_plots.pdf", width = 10, height = 9, dpi = 600)

# Site-level random effects and correlation ---------------------------
site_effects <- interaction_model_t %>%
  spread_draws(r_site_id[site_id, term]) %>%
  mutate(effect_type = ifelse(term == "Intercept", "intercept", "slope")) %>%
  group_by(site_id, effect_type) %>%
  summarise(
    mean = mean(r_site_id),
    lower = quantile(r_site_id, 0.025),
    upper = quantile(r_site_id, 0.975),
    .groups = "drop"
  ) %>%
  pivot_wider(
    id_cols = site_id,
    names_from = effect_type,
    values_from = c(mean, lower, upper),
    names_glue = "{effect_type}_{.value}"
  )

# get fixed effects
fixed_eff <- (fixef(interaction_model_t))

fixed_eff

global_trend <- fixed_eff["centered_year", "Estimate"]

# get the INTERACTION terms (slope differences by location)
# these are the ones with "centered_year:location" in the name
interaction_names <- grep("centered_year:location", rownames(fixed_eff), value = TRUE)
location_slope_effects <- fixed_eff[interaction_names, "Estimate"]

# clean up names (remove "centered_year:" prefix)
names(location_slope_effects) <- gsub("centered_year:location", "", interaction_names)

print(location_slope_effects)  # Should show SSI, EI, SOI

# Add location info to site_effects 
if(!"location" %in% colnames(site_effects)) {
  site_effects <- site_effects %>%
    left_join(model_data_clean %>% 
                select(site_id, location) %>% 
                distinct(), 
              by = "site_id")
}

# Calculate total slope for each site
site_effects <- site_effects %>%
  mutate(
    #  location slope effects for this site (0 for reference location, likely "AP")
    location_slope_effect = case_when(
      location == "SSI" ~ location_slope_effects["SSI"],
      location == "EI" ~ location_slope_effects["EI"],
      location == "SOI" ~ location_slope_effects["SOI"],
      location == "SSW" ~ 0,  # Not in model? Check if exists
      location == "SGI" ~ 0,  # Not in model? Check if exists
      TRUE ~ 0  # Reference location (AP)
    ),
    total_slope = global_trend + location_slope_effect + slope_mean,
    pct_change_3gen = (exp(total_slope * 3 * GENERATION_LENGTH) - 1) * 100
  )

# check one colony to evaluate if worked

site_effects %>% filter(site_id == "BAIL") %>%
  select(site_id, location, slope_mean, location_slope_effect, total_slope, pct_change_3gen)

# check BAIL (should be in SSI)
data.frame(site_effects %>% filter(site_id == "BAIL") )


# add size category and colony size info
colony_info <- model_data_clean %>%
  group_by(site_id) %>%
   
  summarise(
    first_year = min(season_starting, na.rm = TRUE),
    last_year = max(season_starting, na.rm = TRUE),
    first_count = first(penguin_count[season_starting == first_year]),
    first_accuracy = first(accuracy[season_starting == first_year]),
    last_count = first(penguin_count[season_starting == last_year]),
    last_accuracy = first(accuracy[season_starting == last_year]),
    time_interval = last_year - first_year,
    total_change = ifelse(first_count > 0, ((last_count / first_count) - 1) * 100, NA),
    annual_change = total_change / time_interval,
    mean_count = mean(penguin_count, na.rm = TRUE),
    n_years = n_distinct(season_starting),
    lon=mean(lon),
    lat=mean(lat),
    .groups = 'drop'
  ) %>%
  mutate(total_pop = sum(mean_count),
         weight = mean_count / total_pop)
  

site_effects <- site_effects %>%
  left_join(colony_info, by = "site_id")

head(site_effects)

# Extract correlation between intercept and slope
vc <- VarCorr(interaction_model_t)
vc$accuracy_factor
vc$site_id$cor

ranef_summary <- summary(interaction_model_t)$random$site_id
cor_est <- ranef_summary["cor(Intercept,centered_year)", "Estimate"]
cor_lower <- ranef_summary["cor(Intercept,centered_year)", "l-95% CI"]
cor_upper <- ranef_summary["cor(Intercept,centered_year)", "u-95% CI"]


# ---------- site effects geographically ---------------


library(sf)
library(dplyr)
library(terra)

# 

land<-st_as_sf(vect("C:/GIS/add_coastline_medium_res_polygon_v7_10.shp/add_coastline_medium_res_polygon_v7_10.shp")
)
land<-st_transform(land,4326)
plot(land)


FAO<-st_as_sf(vect("C:/GIS/FAO_Major_FIshing_Areas/FAO_48_subareas.shp"))%>%
  st_transform(4326)

plot(FAO)

locations<-st_as_sf(vect("C:/GIS/FAO_Major_FIshing_Areas/FAO_48_1.shp"))%>%
  st_transform(4326)
plot(locations)

table(locations$F_DIVISION)

table(FAO$F_SUBAREA)

site_effects<-site_effects%>%
  mutate(magnitude=abs(slope_mean),
         trend=ifelse(slope_mean<0,"-","+"))

data.frame(site_effects%>%
             filter(site_id=="BAIL"))

model_data_clean%>%
  filter(site_id=="BAIL")


model_data_clean %>%
  filter(site_id=="BAIL")%>%
  ggplot(aes(season_starting,penguin_count))+geom_point()

# create the two plots separately
mappt <- ggplot() +
  geom_sf(data = FAO, fill = NA, colour = "grey30") +
  geom_sf(data = locations, fill = NA, colour = "steelblue", linewidth = 1) +
  geom_sf(data = land, fill = "grey85", colour = "grey40") +
  geom_point(data = site_effects, 
             aes(x = lon, y = lat, 
                 size = mean_count,
                 fill = total_slope,
                 shape = ifelse(total_slope < 0, "-", "+")),
             alpha = 0.85) +
  scale_fill_gradient2(high = "blue", mid = "grey90", low = "red3",
                       midpoint = 0, name = "Slope") +
  scale_size_continuous(range = c(1, 10), name = "Colony size") +
  scale_shape_manual(values = c("+" = 21, "-" = 25), name = "Trend") +
  coord_sf(xlim = c(-70, -42), ylim = c(-68, -60)) +
  theme_bw() +theme(legend.position = "bottom")+
  ggtitle(label = "a. ")

mappt


boxpt <- ggplot(site_effects, aes(y=total_slope)) +
  geom_hline(yintercept = 0, linetype = "dashed") +
  geom_boxplot(width=0.5,fill="grey50") + 
  xlim(-1,1)+
  theme_bw() +
  xlab("Location") +
  ylab("Slope") +
  ggtitle(label = "b.")+
  theme(axis.text.x = element_blank(),
        axis.text.y = element_text(size=8))+
  #ylim(-0.25,0.075)+
  facet_wrap(location~.,nrow=1)

boxpt

combined_inset <- mappt + 
  inset_element(boxpt, left = 0.7, bottom = 0.01, right = 0.98, top = 0.4)

print(combined_inset)

ggsave("Results/colonies_trends.pdf", width = 8, height = 7, dpi = 600)


#---------- risk tables----------------

colony_info <- model_data_clean %>%
  group_by(site_id,location) %>%
  
  
  summarise(
    first_year = min(season_starting, na.rm = TRUE),
    last_year = max(season_starting, na.rm = TRUE),
    first_count = first(penguin_count[season_starting == first_year]),
    first_accuracy = first(accuracy[season_starting == first_year]),
    last_count = first(penguin_count[season_starting == last_year]),
    last_accuracy = first(accuracy[season_starting == last_year]),
    time_interval = last_year - first_year,
    total_change = ifelse(first_count > 0, ((last_count / first_count) - 1) * 100, NA),
    annual_change = total_change / time_interval,
    mean_count = mean(penguin_count, na.rm = TRUE),
    n_years = n_distinct(season_starting),
    lon=mean(lon),
    lat=mean(lat),
    .groups = 'drop'
  ) %>%
  mutate(total_pop = sum(mean_count),
         weight = mean_count / total_pop)


# derive percent change over 3 generations (28.2 years) -------------
# For each site, compute total slope = b_centered_year + interaction + r_site_id
population_draws <- interaction_model_t %>%
  spread_draws(b_centered_year,
               `b_centered_year:locationSSI`,
               `b_centered_year:locationEI`,
               `b_centered_year:locationSOI`,

               
               r_site_id[site_id, term]) %>%
  filter(term == "centered_year") %>%
  left_join(colony_info, by = "site_id") %>%
  mutate(
    site_slope = case_when(
      location == "AP"     ~ b_centered_year + r_site_id,
      location == "SSI"    ~ b_centered_year + `b_centered_year:locationSSI` + r_site_id,
      location == "SOI"  ~ b_centered_year + `b_centered_year:locationSOI` + r_site_id,
      location == "EI"    ~ b_centered_year + `b_centered_year:locationEI` + r_site_id,

      TRUE ~ b_centered_year + r_site_id
    )
  )


# Population-weighted metrics
population_risk <- population_draws %>%
  group_by(.draw) %>%
  summarise(
    weighted_slope = sum(site_slope * weight, na.rm = TRUE),
    pop_pct_change = (exp(weighted_slope * 3 * GENERATION_LENGTH) - 1) * 100,
    total_increase = sum(mean_count[site_slope > 0] * (exp(site_slope[site_slope > 0] * 3 * GENERATION_LENGTH) - 1), na.rm = TRUE),
    total_decline = sum(abs(mean_count[site_slope < 0] * (exp(site_slope[site_slope < 0] * 3 * GENERATION_LENGTH) - 1)), na.rm = TRUE),
    .groups = "drop"
  ) %>%
  mutate(
    net_change = total_increase - total_decline,
    compensation_ratio = total_increase / total_decline
  )


# summarize population-level risk ------------------------------------
n_draws <- nrow(population_risk)

pop_summary <- population_risk %>%
  summarise(
    median_pop_change = median(pop_pct_change),
    median_net_change= median(net_change),
    lower_pop_change = quantile(pop_pct_change, 0.025),
    upper_pop_change = quantile(pop_pct_change, 0.975),
    
    median_net_change = median(net_change),
    net_change_lower = quantile(net_change, 0.025),
    net_change_upper = quantile(net_change, 0.975),
    
    n_risk_30 = sum(pop_pct_change <= -30),
    n_risk_50 = sum(pop_pct_change <= -50),

    n_compensation = sum(net_change > 0)
  ) %>%
  mutate(
    risk_30pct = n_risk_30 / n_draws,
    risk_30_lower = qbeta(0.025, n_risk_30, n_draws - n_risk_30 + 1),
    risk_30_upper = qbeta(0.975, n_risk_30 + 1, n_draws - n_risk_30),
    risk_50pct = n_risk_50 / n_draws,
    risk_50_lower = qbeta(0.025, n_risk_50, n_draws - n_risk_50 + 1),
    risk_50_upper = qbeta(0.975, n_risk_50 + 1, n_draws - n_risk_50),
    prob_compensation = n_compensation / n_draws,
    comp_lower = qbeta(0.025, n_compensation, n_draws - n_compensation + 1),
    comp_upper = qbeta(0.975, n_compensation + 1, n_draws - n_compensation)
  )

print(pop_summary)

# final risk table --------------------------------------------
risk_table <- data.frame(
  Metric = c("Population trend (% change)",
             "Risk of ≥30% decline",
             "Risk of ≥50% decline",
             "Probability of compensation",
             "Net population change (nests)"),
  Estimate = c(
    paste0(round(pop_summary$median_pop_change, 1), "% [",
           round(pop_summary$lower_pop_change, 1), " to ",
           round(pop_summary$upper_pop_change, 1), "]"),
    paste0(round(pop_summary$risk_30pct * 100, 1), "% [",
           round(pop_summary$risk_30_lower * 100, 1), " to ",
           round(pop_summary$risk_30_upper * 100, 1), "]"),
    paste0(round(pop_summary$risk_50pct * 100, 1), "% [",
           round(pop_summary$risk_50_lower * 100, 1), " to ",
           round(pop_summary$risk_50_upper * 100, 1), "]"),
    paste0(round(pop_summary$prob_compensation * 100, 1), "% [",
           round(pop_summary$comp_lower * 100, 1), " to ",
           round(pop_summary$comp_upper * 100, 1), "]"),
    paste0(round(pop_summary$median_net_change, 0), " [",
           round(pop_summary$net_change_lower, 0), " to ",
           round(pop_summary$net_change_upper, 0), "]")
  )
)

print(risk_table)

write.csv(risk_table,"Results/risk_table_D1.csv")

####------- region and risk ------------

head(population_draws)


# compute size_risk from population_draws
region_risk <- population_draws %>%
  group_by(location, .draw) %>%
  summarise(
    category_pct = (exp(mean(site_slope) * 3 * GENERATION_LENGTH) - 1) * 100,
    category_weight = sum(weight),
    .groups = "drop"
  ) %>%
  group_by(location) %>%
  summarise(
    median_pct = median(category_pct),
    lower_pct = quantile(category_pct, 0.025),
    upper_pct = quantile(category_pct, 0.975),
    risk_30 = mean(category_pct <= -30),
    risk_50 = mean(category_pct <= -50),
    contribution = mean(category_weight),
    .groups = "drop"
  )


# crate the table
risk_table_region <- region_risk %>%
  mutate(
    `Population share` = paste0(round(contribution * 100, 1), "%"),
    `Median % change [95% CI]` = paste0(
      round(median_pct, 1), "% [",
      round(lower_pct, 1), "% to ",
      round(upper_pct, 1), "%]"
    ),
    `Risk of ≥30% decline` = paste0(round(risk_30 * 100, 1), "%"),
    `Risk of ≥50% decline` = paste0(round(risk_50 * 100, 1), "%")
  ) %>%
  select(location, `Population share`, `Median % change [95% CI]`,
         `Risk of ≥30% decline`, `Risk of ≥50% decline`)


print(risk_table_region)

# save as csv
write.csv(risk_table_region, "Results/location_risk_table.csv", row.names = FALSE)


# -------- gap map--------------

summary(site_effects)

table(site_effects$location)

head(model_data_clean)

sites<-model_data_clean%>%
  group_by(site_id,size_category)%>%
  summarise(lat=mean(lat))%>%
  select(site_id,size_category)

# create the grouped summary
site_summary <- site_effects%>%
  mutate(n_years_group = cut(n_years, 
                             breaks = c(0, 5, 10, 20, 30, 40, 60),
                             labels = c("0-5", "5-10", "10-20", "20-30", "30-40", "40+"),
                             right = FALSE)) %>%
  group_by(n_years_group,location) %>%
  summarise(
    median_slope = median(total_slope, na.rm = TRUE),
    q1_slope = quantile(total_slope, 0.1, na.rm = TRUE),
    q3_slope = quantile(total_slope, 0.9, na.rm = TRUE),
    median_col = median(mean_count, na.rm = TRUE),
    q1_col = quantile(mean_count, 0.1, na.rm = TRUE),
    q3_col = quantile(mean_count, 0.9, na.rm = TRUE),
    n_sites = n(),
    n_years=mean(n_years),
    median_lat=mean(lat),
    q1_lat = quantile(lat, 0.1, na.rm = TRUE),
    q3_lat = quantile(lat, 0.9, na.rm = TRUE),
    .groups = 'drop'
  )


# get y-position (adjust multiplier for  log scale)
y_max <- max(model_data_clean$penguin_count, na.rm = TRUE) * 1.5

count_data<-count_data%>%
  mutate(counts_per_col=round((counts/colonies),1))


(ggplot(model_data_clean, aes(size_category, penguin_count, fill = location)) +
  geom_boxplot() +
  scale_y_log10(labels = scales::comma_format()) +
  scale_fill_manual(values = c("steelblue", "red3", "orange3", "grey")) +
  theme_bw() +
  # Add counts label
  geom_text(data = count_data, 
            aes(x = size_category, y = y_max*2, 
                label = paste0("", counts_per_col), group = location),
            position = position_dodge(width = 0.75), 
            size = 2.5) +
  # Add colonies label (positioned slightly lower)
  geom_text(data = count_data, 
            aes(x = size_category, y = y_max , 
                label = paste0("", colonies), group = location),
            position = position_dodge(width = 0.75), 
            size = 2.5)+
    ggtitle(label="a.")+xlab("Colony size category")+ylab("Colony size (nest counts)")+
    theme(legend.position = "inside",
          legend.position.inside = c(0.9,0.3) ))/


(ggplot() +
  # Individual site points (background)
  geom_point(data = site_effects, 
             aes(x = lat, y = total_slope,colour=location), 
             alpha = 0.5, 
             size = 1.5) +


  # Centroid points with error bars
  geom_errorbarh(data = site_summary,
                 aes(xmin = q1_lat, xmax = q3_lat, y = median_slope,
                     group = n_years_group),
                 height = 0.005, linewidth = 1, alpha = 0.7 )+
  geom_errorbar(data = site_summary,
                aes(x = median_lat, ymin = q1_slope, ymax = q3_slope,
                    group = n_years_group),
                width = 0, linewidth = 1, alpha = 0.7) +
  geom_point(data = site_summary,
             aes(x = median_lat, y = median_slope, size = median_col,fill=n_years_group),
             shape = 21,stroke = 1.5) +

  # Color scale for individual points
  geom_smooth(data = site_summary, 
              aes(x = median_lat, y = median_slope), fullrange=T,
              #alpha = 0.75, 
              method = "lm",se=F,linetype="dashed",linewidth=1,colour="black") +
 scale_size_continuous(name = "Mean\nsize", breaks=c(500,2000,4000,8000)) +
 xlim(-66,-60.5)+
  # Reference lines
  geom_hline(yintercept = 0, linetype = "dashed", alpha = 0.5) +
  #geom_vline(xintercept = c(5000,25000), linetype = "dashed", alpha = 0.5) +
  # Labels
  labs(x = "Latitude", 
       y = "Site-level trend",
       title = "b. "
  ) +
  theme_bw() +
  theme(plot.title = element_text(hjust = 0, face = "bold"),
        legend.position = "right")+
  scale_colour_manual(values=c("steelblue","red3","orange3","grey"))+
  scale_fill_manual(values=c("steelblue","green3","yellow3","orange2","red3"),name = "N years")+
  theme(legend.position = "inside",
        legend.position.inside = c(0.2,0.2) ,
 legend.box="horizontal",
   legend.key.size = unit(0.3, "cm"),      # Size of the color/symbol boxes
 legend.text = element_text(size = 8),   # Text size (smaller than default)
 legend.title = element_text(size = 9),  # Title text size
 legend.spacing = unit(0.2, "cm")))


ggsave("Results/gap_priorities.png", width = 8, height = 8, dpi = 600)


# this is the end, beautiful frend, the end.
```
